# Distinguishing coalescent models - which statistics matter most?

**DOI:** 10.1101/679498

**Authors:** Fabian Freund, Arno Siri-Jégousse

## Abstract

Modelling genetic diversity needs an underlying genealogy model. To choose a fitting model based on genetic data, one can perform model selection between classes of genealogical trees, e.g. Kingman’s coalescent with exponential growth or multiple merger coalescents. Such selection can be based on many different statistics measuring genetic diversity. A random forest based Approximate Bayesian Computation is used to disentangle the effects of different statistics on distinguishing between various classes of genealogy models. For the specific question of inferring whether genealogies feature multiple mergers, a new statistic, the minimal observable clade size, is introduced. When combined with classical site frequency based statistics, it reduces classification errors considerably.

## 1. Introduction

Modelling genetic diversity of a sample of present-day individuals is a key tool when reconstructing the evolutionary history of populations or scanning the genome for regions under selection, see e.g. [CB17, Section “Nonequilibrium theory”, p. 1026]. For a genetic region without recombination, a standard approach is to model the genealogical tree and the mutations along its branches as random processes. The genealogy thus is given by a random tree with *n* leaves. On this, mutations are modelled as an independent Poisson process along the edges of the coalescent tree.

Several genealogical tree models have been proposed (see Table 1 for a summary. The most widely used model, Kingman’s *n*-coalescent, is a strictly bifurcating random tree [Kin82]. It approximately describes the genealogy of a fixed-size population under neutral evolution if offspring numbers per parent are not too variable, e.g. if the population is described by a (haploid or diploid) Wright-Fisher model or a Moran model with fixed size *N*, see [Kin82]. More precisely, the *n*-coalescent is the limit (in distribution) of the genealogies in the discrete populations as *N → ∞*, if one treats 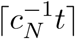 generations as one unit of time in the coalescent tree (which has a continuous time scale). Here, *c*_*N*_ is the coalescence probability, i.e. the probability that two individuals, picked in any generation, have a common parent. Variants of Kingman’s *n*-coalescent as limits of genealogies from Wright-Fisher models with varying population size and/or population structure are available, see e.g. [GT94], [WH98]. These variants still are strictly bifurcating trees. Barring recombination, both haploid and diploid populations are well-captured by Kingman’s *n*-coalescent and its variants.

**Table 1.** List of coalescent classes and their corresponding evolutionary scenario. References are listed within the introduction

| Coalescent model | Type of evolution | Examples |
| --- | --- | --- |
| Kingman's<br>$n$-coalescent | Wright-Fisher model<br>Moran model | Populations with fixed size<br>of mammals, not under<br>strong selection pressure |
| Beta-$n$-coalescent | Sampling constant size<br>population from i.i.d. power-<br>law offspring distributions<br>per individual | Haploid species w. sweepstake<br>reproduction, virus<br>populations w.<br>super spreaders |
| Dirac $n$-coalescent | Moran model w. occasionally<br>single very large family<br>(<u>fixed</u> fraction $p$ of population) | (Haploid) sweepstake<br>reproduction w. fixed<br>sweepstake size |
| Beta-Ξ-$n$-coalescent | As Beta-$n$-coalescent, but off-<br>spring per randomly<br>mating pair of diploids | Diploid species w.<br>sweepstake reproduction |

**Table 2.**
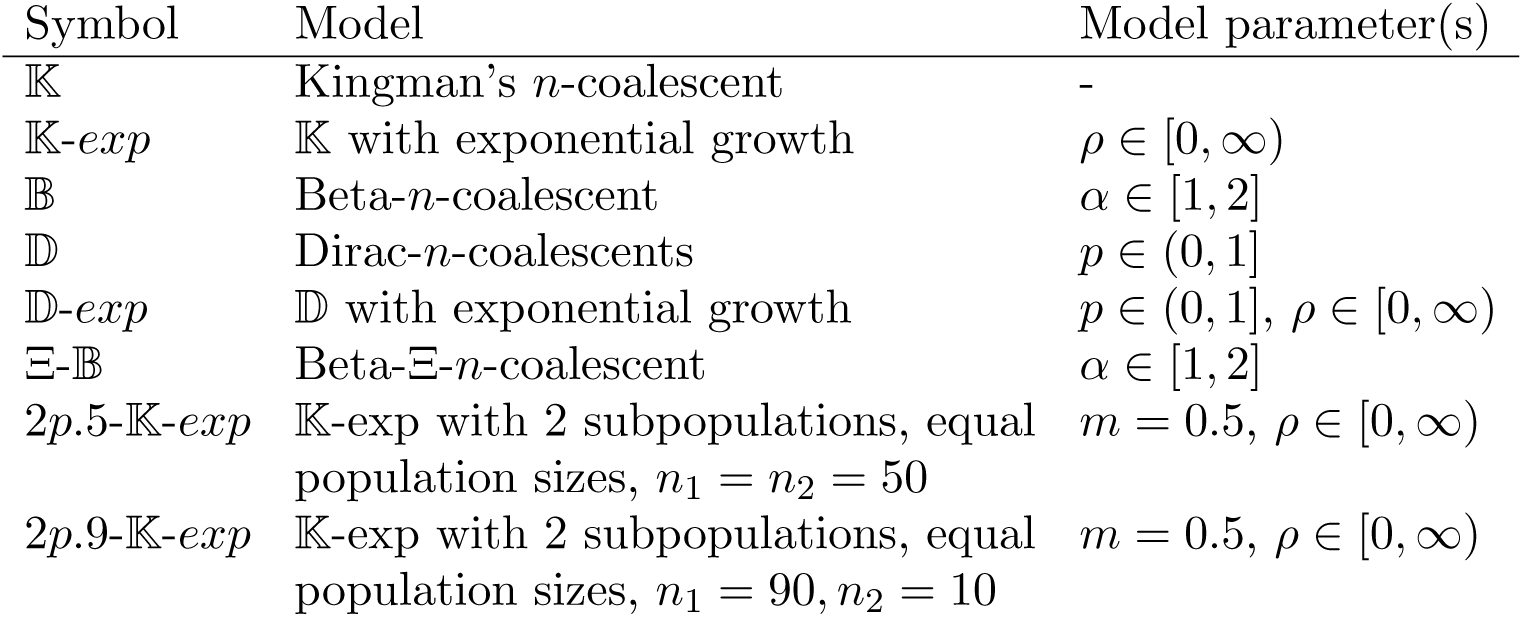
Coalescent classes and parameters ranges

However, some populations will feature genealogies not well approximated by Kingman’s *n*-coalescent and its bifucating variants. Certain modes of rapid selection, affecting the whole genome, should lead to the Bolthausen-Sznitman *n*-coalescent, e.g. see [NH13], [DWF13], [Sch17]. This genealogy model likely produces multifurcating genealogical trees. Even a succession of hard selective sweeps may cause a multifurcating genealogy (though not the same model), see [DS05].

Generally, if one considers the genealogy in an arbitrary Cannings model, a fixed-size discrete-time population model where offspring sizes are exchangeable among parents, a whole class of multifurcating trees, the Λ- or Ξ-*n*-coalescents, emerges as possible weak limits of the genealogies for *N → ∞*, again with the analogous rescaling as for Kingman’s *n*-coalescent, see [MS01]. Non-Kingman *n*-coalescents arise if the distribution of offspring size per parent is variable enough, see [Möh98]. They have been shown to explain the genetic diversity of several maritime species better than Kingman’s *n*-coalescent and its variants, see e.g. [EW06], [NNY16] or [HSJB19]. They seem to capture well the concept of sweepstake reproduction, where one individual may produce a considerable part of the population by chance. Two specific haploid models proposed for this are leading to the Eldon-Wakeley *n*-coalescents, see [EW06], and to the Beta-*n*-coalescents, see [Sch03]. However, for diploid populations, the multiple merger mechanism is modified from its haploid version, see [BLS18]. Other mechanisms may also lead to these coalescent limits, e.g. [HP19] argues that Beta-*n*-coalescents may describe genealogies in viral populations where super-spreaders tend to infect a large number of individuals. Again, variants for varying population sizes in the pre-limit models are available, see [MHAJ18], [Fre20], [KWB19], [GCMPSJ19]. For further discussion where multiple merger genealogies may arise, see the reviews [TL14] and [ILM^+^16].

How can one distinguish these genealogical models based on a sample of *n* genomic sequences? Available methods using the full sequence information do not scale well with data sets with many individuals and/or regions with many mutations [SBB13], [Ste09], [Kos18]. Thus, less costly methods based on a summary statistic of the genomic information, the site-frequency spectrum (SFS), have been proposed, see e.g. [EBBF15], [Kos18]. The SFS records how many mutations appear exactly *i* times in the sample for each *i ∈*{1, …, *n*}. These methods are noisy for one or few regions, but have fairly low error rates for data taken across the genome for many unlinked loci, see [Kos18]. However, even whole genomes may not contain many unlinked loci. For instance, some bacterial genomes consist of a single chromosome with neglectable recombination, e.g. in *Mycobacterium tuberculosis*, also several fungi have comparably small genomes with large linkage blocks. For cancer cells, (one-locus) Beta-*n*-coalescents with exponential growth have been proposed as a genealogy model for copy numbers (specific mutations) in cancer cells [KVS^+^17] within multiple regions. In this study, quantiles of the SFS and several additional statistics were used as summary statistics for inference based on Approximate Bayesian Computation (ABC). Low classification errors (≤2%) were reported, which were much lower than the SFS-based classification errors from [EBBF15]. The discrepance of error rates may come from slightly different hypotheses, a different approach to mutation rates, a different ABC approach and/or the use of additional summary statistics.

Motivated by this drop in error rates, our goal is to investigate which statistics are best suited to distinguish different classes of *n*-coalescents. Additionally to the statistics from [KVS^+^17], we consider some further common statistics as well as a new statistic, the smallest allele frequency among non-private mutations observed in one individual.

These statistics are used in an ABC framework. A simulation-based method is a reasonable choice since the distributional characteristics of the diversity statistics used are not fully understood, see e.g. [BB19, Sect. 2.2.2]. We use a slight modification of the ABC approach based on random forests from [PME^+^15] that performs well with many, potentially uninformative statistics and that measures the importance of each statistic to distinguish between the hypotheses.

### 1.1. Methods

The genealogical trees are all given by Λ- or Ξ-*n*-coalescents, which are random trees with a Markovian structure. We start with *n* leaves at time 0, which start *n* branches, and build the tree until its root is reached. The branches represent (ancestral) lineages of a sample of *n* individuals. If *b* branches are present at a time *t, k* of these, chosen at random, will be merged with rate *λ*_*b,k*_. In the case of Λ-*n*-coalescents, *k* lineages are merged into one new lineage, in other words they are joined in a node of the tree and a new branch towards the root is started. These rates are characterized by

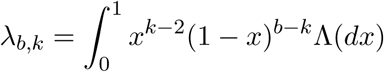

with a finite measure Λ on [0, 1], see [Pit99]. For Ξ-*n*-coalescents, *k* = (*k*_1_, …, *k*_*m*_) sets of lineages are each merged into one new lineage, so *m* new nodes are added from which *m* new branches extend (the new node *i* has thus in-degree 1 and outdegree *k*_*i*_). The rate 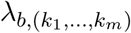 of such a merger can be expressed in terms of a finite measure Ξ on the simplex, as introduced in [Sch00].

As introduced above, all *n*-coalescents can be seen as limit trees of genealogies of a sample size *n* in discrete populations of fixed size *N → ∞* across all generations. If population size may change, but stays of order *N* across generations, the genealogy tree is then given by a *n*-coalescent characterized by the same measure as in the fixed size case, but with time changed by a deterministic function, see [GT94], [MHAJ18], [Fre20], [KWB19]. As Λ-coalescent model classes, we consider Kingman’s *n*-coalescent, Dirac *n*-coalescents with Λ = *δ*_*p*_ for *p ∈* (0, 1], which are a subclass of Eldon-Wakeley *n*-coalescents, and Beta-*n*-coalescents with Λ = *Beta*(2 −*α, α*) for 1 ≤ *α* ≤ 2. For the first two classes, we also consider the coalescent models within exponentially growing populations with growth rate *ρ*, which is reflected by a time change *t ↦* (*γρ*)^−1^ (*e*^*γρt*^ *–* 1) of the coalescent model. The parameter *γ* is determined by the underlying discrete population model, see [Fre20]. For Kingman’s *n*-coalescent, we use *γ* = 1, which corresponds to seeing Kingman’s *n*-coalescent as the limit genealogy of genealogies in Wright-Fisher models, see [GT94]. For the Dirac *n*-coalescent, *γ* reflects the rarity of large families in the discrete population model, see [MHAJ18], we choose *γ* = 1.5. We also consider one class of (non-Λ) Ξ-*n*-coalescents, the Beta-Ξ-*n*-coalescents from [BCEH16] and [BLS18]. To assess population structure as a confounding factor, we also consider the structured (Kingman’s) *n*-coalescent with exponential growth. We assume 2 subpopulations with equal size, symmetric migration with scaled migration rate *m* (in units of 4*N*) and samples of sizes *n*_1_, *n*_2_ from each subpopulation summing up to *n*. See Table 1 for the short names for each coalescent model class and its (theoretical) parameter ranges.

### 1.2. Simulation of coalescent models

We employ different prior parameter distributions for coalescent model parameters and mutation rates.

To be consistent with [KVS^+^17], for a scenario comparable to their setup, denoted by Scenario 1, we use a predefined mesh of mutation rates *θ* and coalescent parameters. The mutation rates are identical for all model classes considered (and each parameter choice therein).

However, the distribution of total length of coalescent processes differs strongly between (and within) model classes, and thus does the total number of mutations. If statistics are strongly affected by the mutation rate, this may either lead to a bad fit to the observed genetic diversity (since the parameter range used simply produces too many or too few mutations for specific models), or parameter ranges have to be very large to counteract these effects. In our case, most statistics are relatively sensitive to changes in the mutation rate. To make models comparable, we start with setting the mutation rate in each model such that the expected number of mutations observed is identical. In other words, we assume, as e.g. in [EBBF15], an (integer) number *s* of observed mutations and set *θ* as the generalized Watterson estimator 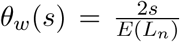, where *L*_*n*_ is the total branch length in the specific model we are looking at. Choosing different coalescent parameters will cause different *θ* values within one model class. Naturally, we do not know the true mutation rate, so we should allow some fluctuation of *θ* for each model. We do this either by picking the number *s* of observed mutations uniformly from a set of integers, or by ‘blurring’ *θ* by drawing the mutation rate from [*b*^−1^*θ*_*w*_(*s*), *bθ*_*w*_(*s*)] for one fixed value of *s* for an integer *b* (we choose 2 or 10). We do the latter by a discrete prior symmetric on a log scale. We first draw *X* from a binomial distribution *B*(*b*, 0.5) and set the mutation rate to *θw*(*s*)*b*^(2*X*−*b*)*/b*^.

Apart from Scenario 1, we employ continuous prior distributions where possible. For the Beta and Dirac *n*-coalescent models, we use the uniform distribution on the parameter range. For Kingman’s *n*-coalescent with exponential growth, we use an uniform prior on the log-scale for the growth parameter *ρ* with an added atom at *ρ* = 0. This means that for a given parameter range [*a, b*] and an atom of size *a*, with probability 1 − *a* we draw *ρ*^*′*^ from the continuous uniform distribution on [*log*(*a*), *log*(*b*)] and set *ρ* = *exp*(*ρ*^*′*^), and with probability *a* we set *ρ* = 0. Computational constraints led us to use discrete priors for the Beta-Ξ-*n*-coalescent (coalescent process rates are expensive to compute), for the Dirac-coalescent with exponential growth (Watterson’s estimator is costly to compute) and for Kingman’s *n*-coalescent with exponential growth and population structure (Watterson estimator is approximated via simulation, which is costly). All discrete priors are discrete uniform distributions on the set of parameters.

See Table 3 for the different prior distributions for coalescent parameters. The specific scenarios for all three choices for mutation rates are shown in Table 3. All scenarios 1-5, consisting of specific choices of *coalescent model × sample size n × mutation model* are simulated twice with 175,000 times with an uniform prior on the model class parameter. For the mutation rate, we either draw the mutation parameter *θ, s* uniformly from a discrete set, or we draw with binomial probabilities from a log-equidistant discrete set around *θ*_*w*_(*s*) (as described above).

**Table 3.**
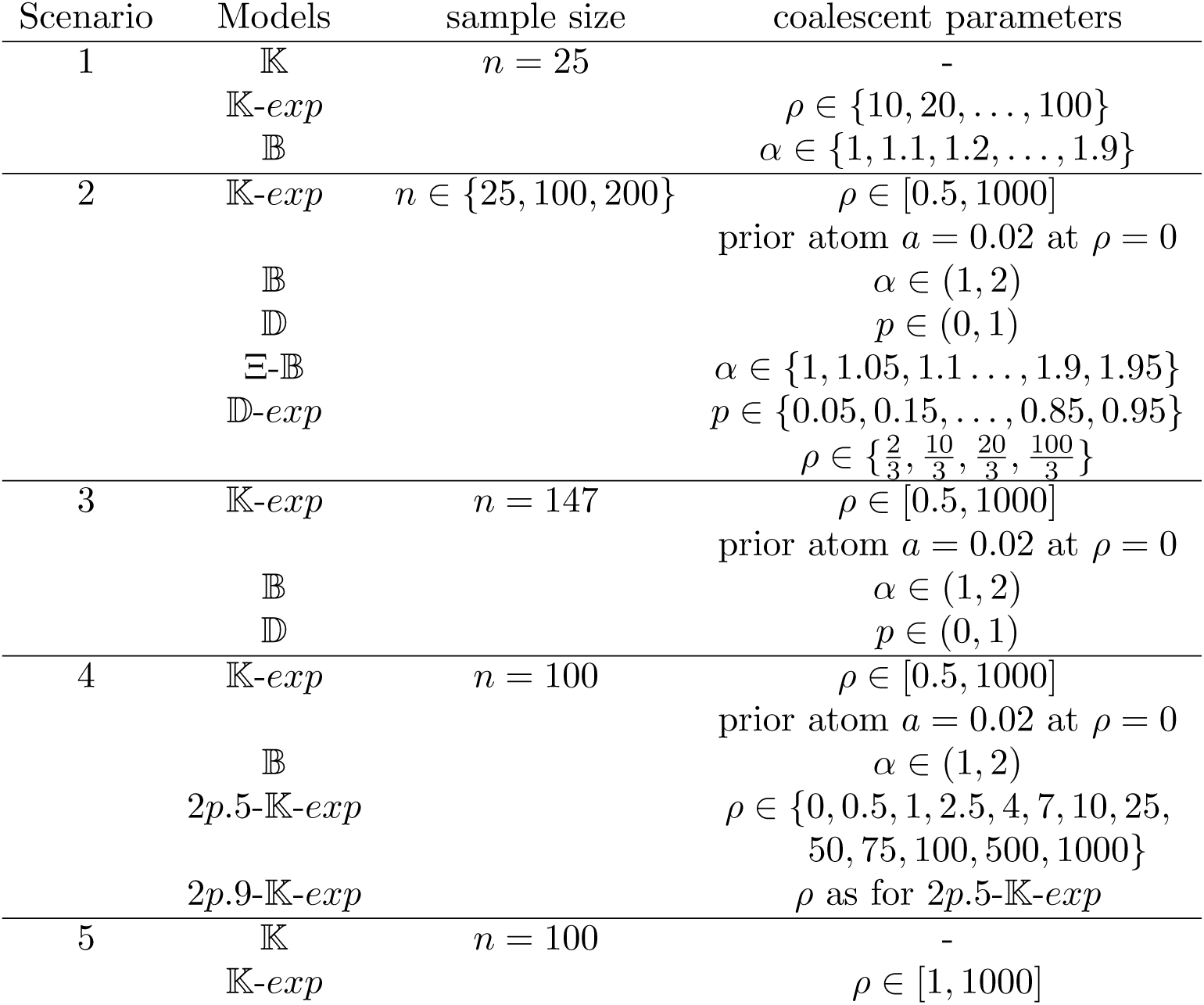
Simulation settings - model classes. Model selection can be both performed for all model classes within a scenario or only for a subset.

**Table 4.** Simulation settings - mutation models

| Scenario | mutation model |
| --- | --- |
| 1 | $\theta \in \{10, 20, \dots, 100\}$ |
| 2 | $s \in \{15, 20, 30, 40, 60, 75\}$ |
| 3 | $\theta \in [10^{-1}\hat{\theta}_w(454), 10\hat{\theta}_w(454)]$ |
| 4 | $s = 75$ |
| 5 | $s \in \{100, 150, 200\}$ |

For simulation of 𝕂, 𝕂-*exp*, 2*p*.5-𝕂-*exp* and 2*p*.9-𝕂-*exp*, we perform simulations with Hudson’s `ms`, implemented in the `R` package `phyclust` [Che11]. 𝔻, 𝔹 and Ξ-𝔹 were simulated with `R`, see Section 4 for code availability. 𝔻-*exp* was simulated as described in [MHAJ18], using the implementation available at https://github.com/Matu2083/MultipleMergers-PopulationGrowth. To compute the (generalized) Watterson estimator 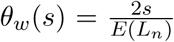, where *s* is the number of observed segregating sites, we employ different methods to compute (resp. approximate) the expected total length *E*(*L*_*n*_) of the genealogy. For 𝕂, we simply compute 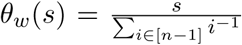. For 𝕂-*exp*, we use a recursive computation approach for *E*(*L*_*n*_) as described in [EBBF15], available from http://page.math.tu-berlin.de/~eldon/programs.html. For 𝔻-*exp*, we again use a recursive approach for computing *E*(*L*_*n*_) as described in [MHAJ18] (url as above). For 𝔹, 𝔻 and Ξ-𝔹, we implemented the recursion for *E*(*L*_*n*_) from [Möh06, Eq. 2.3] in `R` (see Section 4 for availability). For 2*p*.5-𝕂-*exp* and 2*p*.9-𝕂-*exp*, we approximate *E*(*L*_*n*_) by averaging the total length of 10,000 simulated *n*-coalescents for each parameter set (again via `ms`).

### 1.3. Summary statistics

Each simulation produces a matrix with elements in {0, 1} with *n* rows and a variable number *s* of columns. The rows and columns correspond to the number of simulated sequences and to the number of mutations therein. We call it a (simulated) SNP matrix. We use a variety of standard summary statistics from population genetics, mostly as in [KVS^+^17], but also introduce a new statistic. All statistics depend on the coalescent model, *θ* and *n*. For sake of readability, we omit this dependence in the notations.

The standard statistics are all based on the following four sets of statistics Let (*c*_*i,j*_)_*i∈*[*n*],*j∈*[*s*]_ be a SNP matrix.

- The site frequency spectrum (*S*_*i*_)_*i∈*[*n*−1]_ (SFS). The r.v. *S*_*i*_ counts how many columns sum up to *i*, corresponding to how many mutations have a mutant allele count of *i*. These are *n* − 1 statistics.

Alternatively, we can also summarize this information by recording the (quantiles of the mutant) allele frequencies 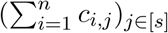.

- The Hamming distances for each pair of rows of the SNP matrix, i.e. the number of pairwise differences between two sequences. These are 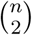 statistics.
- All tree lengths of the phylogenetic tree reconstructed by the neighbor-joining method [SN87], as implemented in the R package `ape` [PS18], based on the set of Hamming distances. These are 2*n* − 2 statistics
- The squared pairwise correlation between sums for each pair of columns, i.e. the pairwise linkage disequilibrium (LD) measure *r*^2^. These are 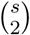 statistics.

We also introduce a set of *n* new statistics, the minimal observable clade size (*O*_*i*_)_*i∈*[*n*]_. Again, let (*c*_*i,j*_)_*i∈*[*n*],*j∈*[*s*]_ be a SNP matrix. Then,

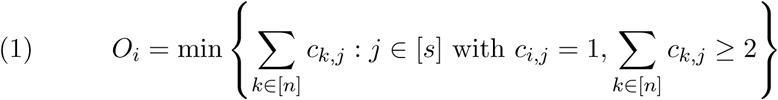

with the convention that *O*_*i*_ = *n* if the condition on *j* is not fulfilled by any *j*. Interpreted in genetic terms, this is the minimal allele count of the mutations of *i* that are shared by at least one other sequence, see Table 5 for an example. We call *O* the minimal observable clade size, since it corresponds to the size of the minimal clade including *i* and at least one other sequence that can be distinguished from the data. See [FSJ19] for further mathematical properties of *O*.

**Table 5.**
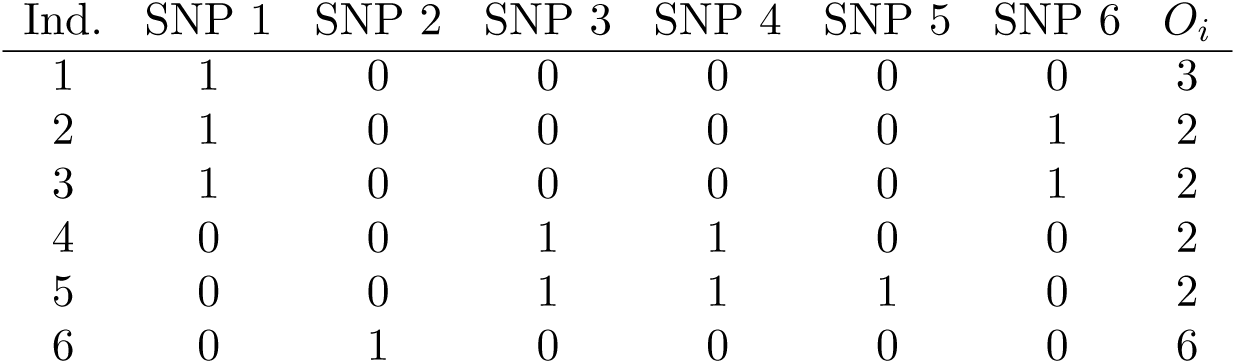
Example computation for the minimal observable clade sizes (*O*_*i*_)_*i∈*[*n*]_. The columns between the vertical lines are a SNP matrix, where 0 denotes the ancestral type and 1 denotes the derived type.

| Ind. | SNP 1 | SNP 2 | SNP 3 | SNP 4 | SNP 5 | SNP 6 | $O_i$ |
| --- | --- | --- | --- | --- | --- | --- | --- |
| 1 | 1 | 0 | 0 | 0 | 0 | 0 | 3 |
| 2 | 1 | 0 | 0 | 0 | 0 | 1 | 2 |
| 3 | 1 | 0 | 0 | 0 | 0 | 1 | 2 |
| 4 | 0 | 0 | 1 | 1 | 0 | 0 | 2 |
| 5 | 0 | 0 | 1 | 1 | 1 | 0 | 2 |
| 6 | 0 | 1 | 0 | 0 | 0 | 0 | 6 |

To reduce the number of statistics, we follow [KVS^+^17] and use both low dimensional summaries of the statistics or quantiles of each of the five sets of statistics. Additionally to quantiles of the different sets of statistics presented above, we use the following summaries of the

- SFS: Tajima’s *D* [Taj89], Fay and Wu’s *H* [FW00], the lumped tail 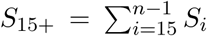, e.g. [Kos18], [EBBF15], and the total number 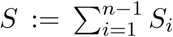 of segregating sites (the number of columns of the_*i*_ SNP matrix). The lumped SFS is (*S*_1_, *S*_2_, …, *S*_15+_)
- Hamming distances: sample mean *π*, i.e. the nucleotide diversity (which can also be computed based on the SFS, see e.g. [Dur08, Example 1.4])
- The sample mean, sample variance and harmonic mean of the observable clade sizes (*O*_1_, …, *O*_*n*_).

We will also use the folded version of the SFS. The folded site frequency spectrum is defined as 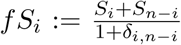 for 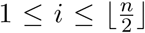, where *δ*_*i,n*−*i*_ is the Kronecker symbol which is 0 unless *i* = *n* − *i* where it equals 1. The lumped fSFS, analogously to the lumped SFS, is (*f S*_1_, *f S*_2_, …, *∑*_*i*≥15_ *f S*_*i*_).

Since our statistics are rather sensitive to changes in the mutation rate, we consider the following statistics that are transformed to be more robust to such changes

- The scaled SFS 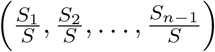 and its lumped version 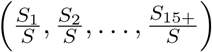
- Quantiles (.1, .3, …, .9) of: the Hamming distances divided by the nucleotide diversity *π*
- Quantiles (.1, .3, …, .9) of 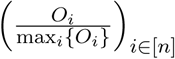

For the reduced sets of statistics mostly used in this study, see Table 6 for short names and setup. For Tajima’s *D* and Fay and Wu’s *H*, we used the function `sfsR` from https://github.com/sdwfrost/popseq. All statistics have been computed with `R`, see Section 4 for code availability.

### 1.4. Model selection via ABC

We perform model selection with the ABC approach introduced by [PME^+^15], based on random forests, which we modify slightly. For this approach, we simulate each model class used with parameters drawn from the prior distribution. Then, bootstrap samples are drawn from the simulations, i.e. we draw with replacement *n*_*trees*_ times a bootstrap sample of size *n*_*boot*_ from *n*_*sim*_ simulations. Then, for each bootstrap sample a decision tree (randomized CART tree) is constructed, whose decision nodes are of the form {*T* > *t*} or {*T* < *t*} for any of the summary statistics used and for *t ∈* ℝ. For each node, a random subset of all statistics is considered and the statistic *T* is picked such that it sorts the bootstrap sample to the model classes as efficient as possible, i.e. so that ideally simulations from different model classes are sorted to different branches at the decision node (low node impurity). In [PME^+^15], this sorting efficiency is measured by the Gini index, which is recommended for model classification. However, we are interested in the effect of different statistics on the model classification and using the Gini index is known to be biased towards picking statistics with larger ranges (more possible values) at decision nodes [SBZH07]. Since we use statistics with different ranges (resp. different numbers of possible values), we account for this bias by correcting for uninformative splits using the approach from [SZ08] as described in [NKW18]. See [NKW18, Sect. 2.3, 2.4] for details.

**Table 6.**
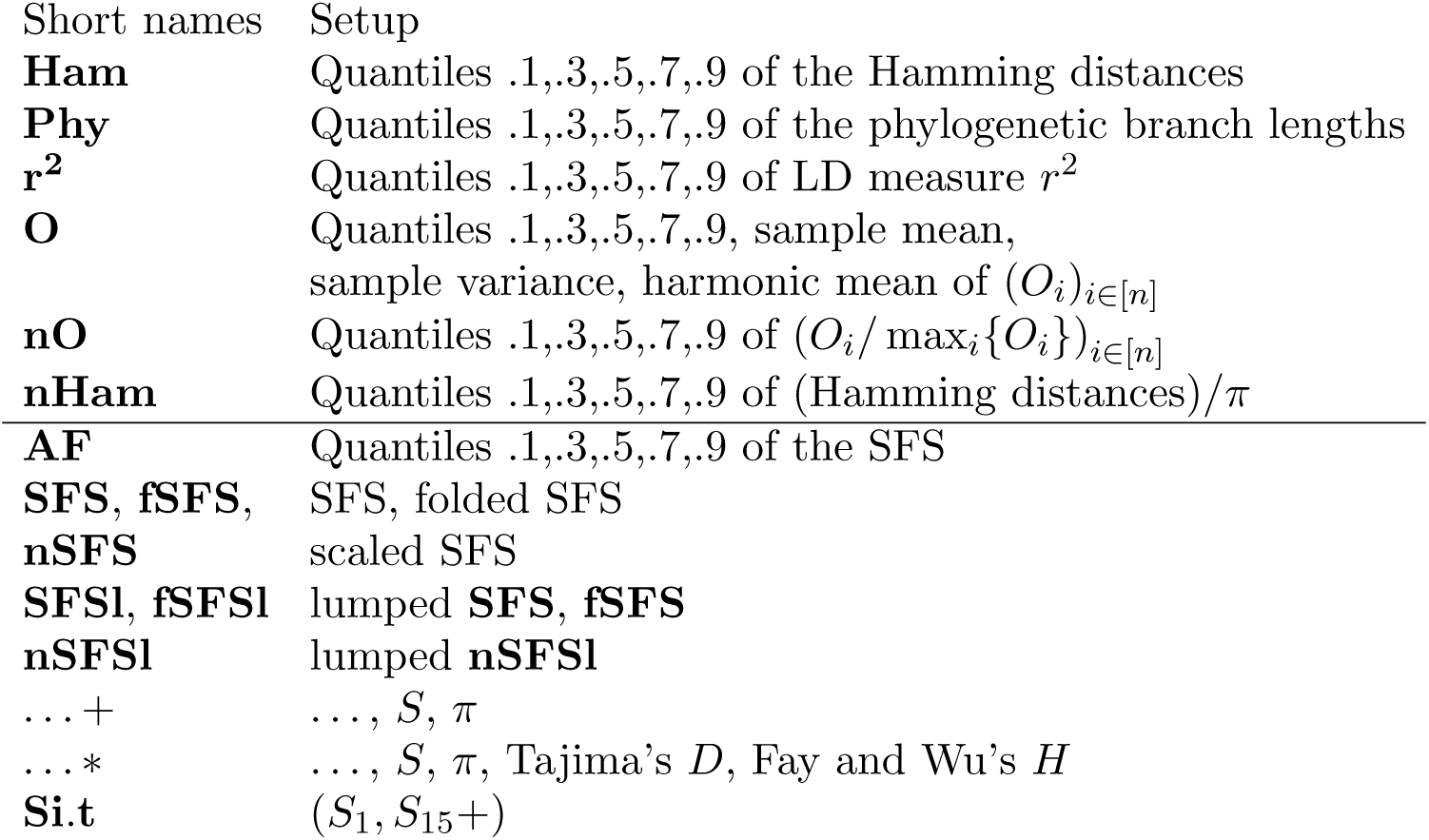
Short names for sets of statistics used for model comparisons. For the sets beneath the line, adding + (*\**) denotes adding the two (four) additional statistics.

Nodes are then added until the bootstrap sample can be perfectly allocated to the model classes. This means sorting the simulations via the decision rules to distinct sets of simulations, so that each set only includes simulations from the same model class *A*. This leaf of the decision tree is then classifying other simulations, resp. real observed data, which are guided to it by the decision tree (also) to model class *A*. The (mis)classification probability for each model class *A*_1_ is estimated by the out-of-bag (OOB) error. For each simulation from a model *A*_1_, the (mis)classification probability can be assessed by recording the proportion of the trees built without this simulation that sort it into model class *A*_1_, *A*_2_, …. The probability of a misclassification (classification error) is then the sum of these proportions who sort the simulation into a different class than the one it comes from. The out-of-bag (classification) error for model class *A*_1_ is given by the mean of the misclassification probabilities over all simulations of *A*_1_. Finally, the mean out-of-bag error is the mean classification error across all model classes.

As an alternative measure for the ability of different statistics to distinguish between model classes, we use variable importance. Variable importance of a statistic *T* is measured as the decrease of node impurity by all nodes using *T* as decision statistic, averaged over the whole random forest (Gini VI measure, with the modification as decribed above).

This ABC approach should cope well with high numbers of potentially uninformative summary statistics (curse of dimensionality) and should perform at least comparable to standard ABC methods. See Section A.5 for a comparison with other ABC methods for Scenario 2. Since we are interested in the variable importances for the presented summary statistics, we omitted the option to include the linear discriminant(s) of the statistics in the ABC analysis, i.e. the linear combination of variables that maximizes difference between model classes for the training data.

For this ABC approach, we used the `R` package `abcrf`. We modified the default use of variable importance as described above in the underlying command `ranger` from the `ranger` package [WZ17].

For any scenario, we perform model selection either between all model classes within the scenario or just for a subset of model classes. For every full set or subset of model classes we consider, we independently perform the following procedure. For each of 2 replications of simulations from the model classes we want to distinguish, we perform 10*×* an ABC model selection with a random forest of *n*_*trees*_ = 500 trees, each constructed from a bootstrap sample with (default) size *n*_*boot*_ = 100, 000. Thus, the mean misclassification probabilities across the 10 runs can also be interpreted as a single run with a random forest of 5,000 trees. We can thus aggrevate both variable importance and mean classification errors by further averaging over the 10 runs. We will report the range across the 10 runs as well as the overall mean. Table 7 provides a common plot legend for all ABC result plots. For code availability, see Section 4.

**Table 7.** Common legend for figures showing classification errors and/or variable importances

|  |  |
| --- | --- |
| Black bar | Range of mean misclassification probabilities<br>(resp. variable importances) over 10 ABC runs<br>in the first set of simulations |
| Gray bar | Range of mean misclassification probabilities<br>(resp. variable importances) over 10 ABC runs<br>in the second set of simulations |
| Asterisk<br>(on a range bar) | shows the mean of all objects that contribute<br>to the range bar |

## 2. Results

We assessed the effects of different sets of diversity statistics on the classification errors for distinguishing different coalescent models. We report changes in classification errors, e.g. if we report a decrease of 2% in error, this is an absolute decrease, e.g. from 12% to 10% misidentification chance.

If a statistic’s variable importance is not reported, it showed no variation among simulations and was omitted from the ABC analysis. Results are available as `R` objects, see Section 4.

### 2.1. Scenario 1: Beta vs. Kingman vs. Kingman with exponential growth

In the first scenario, we partially reproduce the approach in [KVS^+^17], ignoring sequencing and model identification errors and only including a subset of models to select from, see Tables 3 and 4. We consider the statistics as in [KVS^+^17], but adding *π* and **O**. Results are shown in Figure 1.

**Figure 1.**
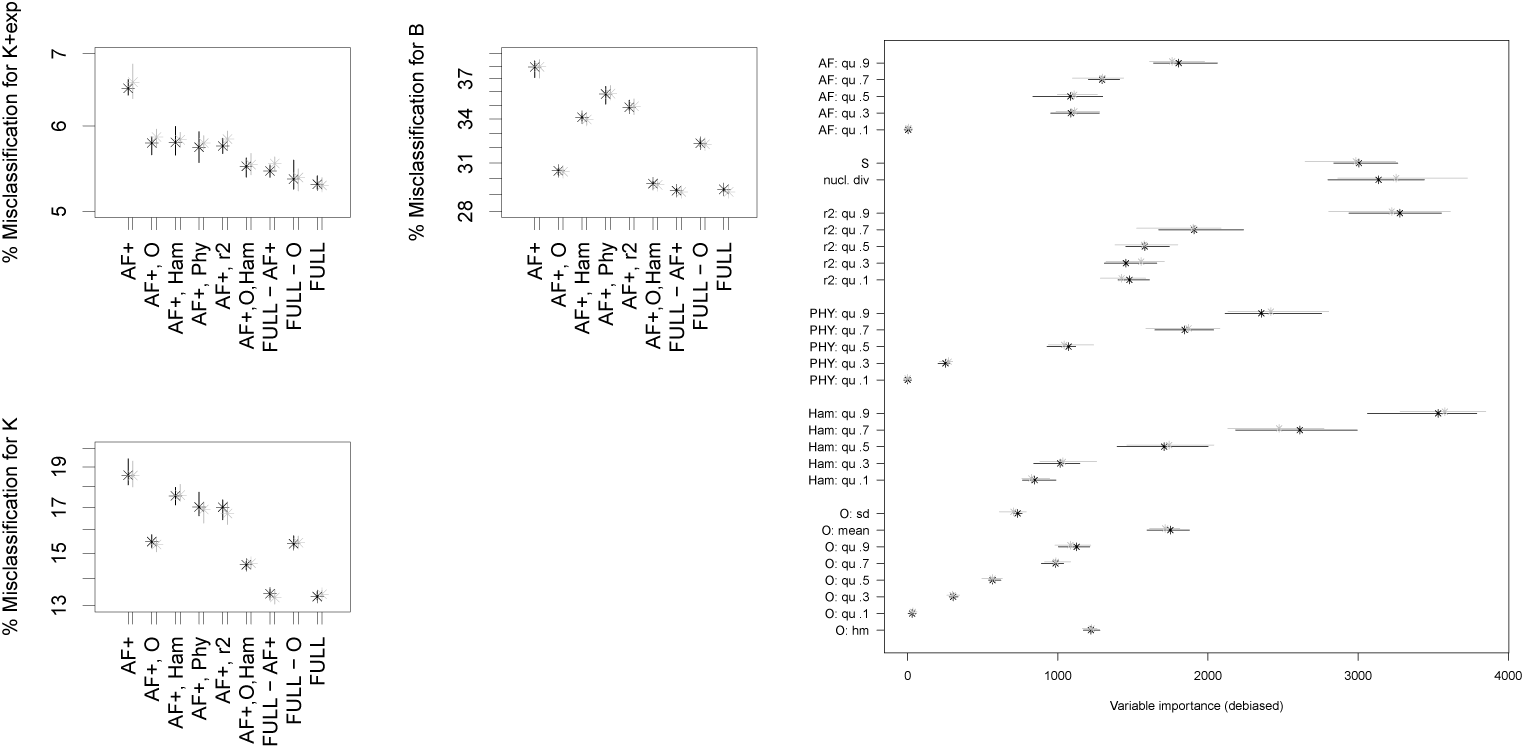
Classification errors (on a log-scale) and variable importances of model comparison between 𝕂-*exp*, 𝕂 and 𝔹 (Scenario 1). The full set of statistics consists of **AF, r**^***2***^, **Phy, Ham, O**, *S* and *π*. For the legend, see Table 7.

We clearly see that the strategy of [KVS^+^17] to add **Phy**, **Ham** and **r**^***2***^ to the (summaries *S, π* and quantiles of) the allele frequencies is beneficial (mean out-of-bag error reduced by > 3% in both replications). Further adding the minimal observable clade sizes leads to even lower classification errors (further mean out-of-bag error reduction by > 1.5%), see Figure 1. The best addition to the site frequency spectrum-based statistics in this scenario is actually **O**, reducing the mean OOB by ≈4%, which is stronger than the effect of **Phy, Ham, r**^***2***^ when added separately. However, the other statistics do feature further information to distinguish the three model classes, reflected by a further reduction in misclassification errors (additional mean OOB reduction of ≥1%) when all statistics are used.

Generally, misidentification is very common for 𝔹, while misidentification for 𝕂 or 𝕂-*exp* is much less common. Classification errors are strongly increased compared to the original study of [KVS^+^17], which we attribute to us classifying only part of their range of parameters in each model (see discussion in Section 3). Scenario 1 uses common mutation rates for each model, leading to quite different expectations of how many segregating sites appear for coalescents with different coalescent parameters (and models). This is due to the differences in total genealogical tree length between models. Thus, the ABC approach should also use this information to select models, i.e. pick statistics for decision nodes that rely on the number of mutations observed. The most important statistics in Scenario 1 are the 0.9 quantiles of the Hamming distances and of *r*^2^, as well as the nucleotide diversity *π* and the number of segregating sites *S*, which are all strongly influenced by the number of mutations observed. Interestingly, no statistic from **O** is among the statistics with highest variable importance. A likely reason, for which will we see more evidence for in Section 5, is that **O** may not be very helpful to distinguish between 𝕂+exp and 𝕂, thus should be chosen as decision statistic only for nodes where sorting between these model classes is not necessary. This should reduce the importance scores.

### 2.2. Scenario 2: Beta vs. Kingman with exponential growth

In Scenario 2, we compare model classes that, on average, produce a similar number of mutations.

#### 2.2.1 *Model selection using* AF+, Ham, O, Phy, r^*2*^

As in Scenario 1, adding statistics to the quantiles of the allele frequency spectrum helps distinguishing the model classes, see Figure 2, and leads to moderate to strong error decrease (mean OOB error reduced by ≥5%). For this scenario, the best addition to the allele frequency quantiles is **O**. For all *n*, it is actually better adding **O** to **AF**+ than adding all the other statistics, and this is more visible for big values of *n* (mean OOB error further reduced by > 1.8% for *n* = 200 if **O** is added). The positive effect of **O** can be slightly further increased by adding **Ham**, the set of statistics with the second largest decrease when added to **AF**+. Adding these two sets of statistics to **AF**+ leads to mean OOB errors very close to those of the full set of statistics. The variable importance scores in Figure 3 show that especially the harmonic mean of **O** is a statistic that distinguishes the model classes well. It is only overtaken for *n* = 25 by the 0.9 quantiles of the allele frequency and *r*^2^. For *n* > 25, further statistics from **O** show high importance.

**Figure 2.**
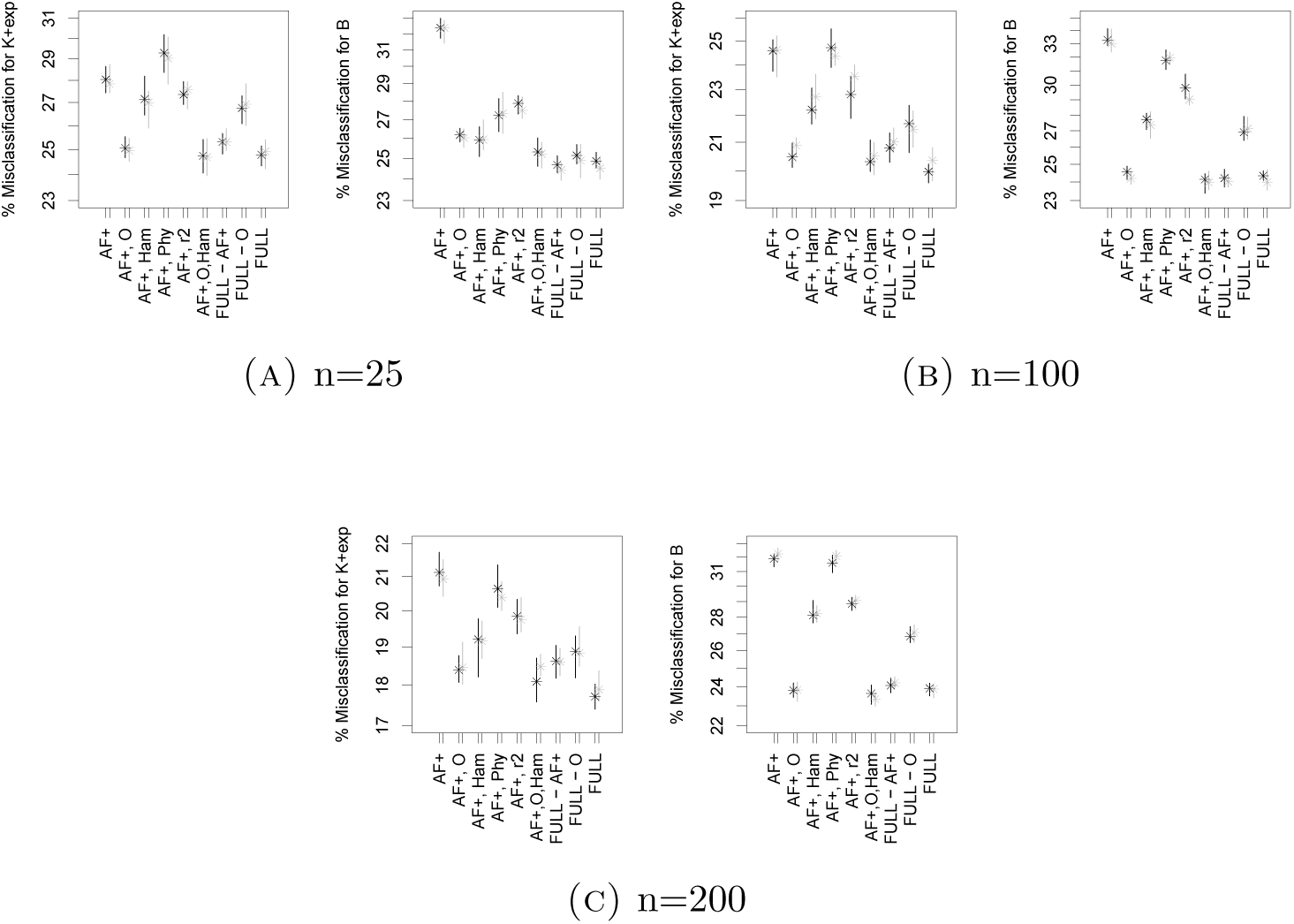
Classification errors (on a log-scale) of model comparison between 𝕂-*exp* and 𝔹 (Scenario 2). See Table 7 for the legend. The full set of statistics consists of **r**^***2***^, **Phy, Ham, O**, *S, π* and **AF**

**Figure 3.**
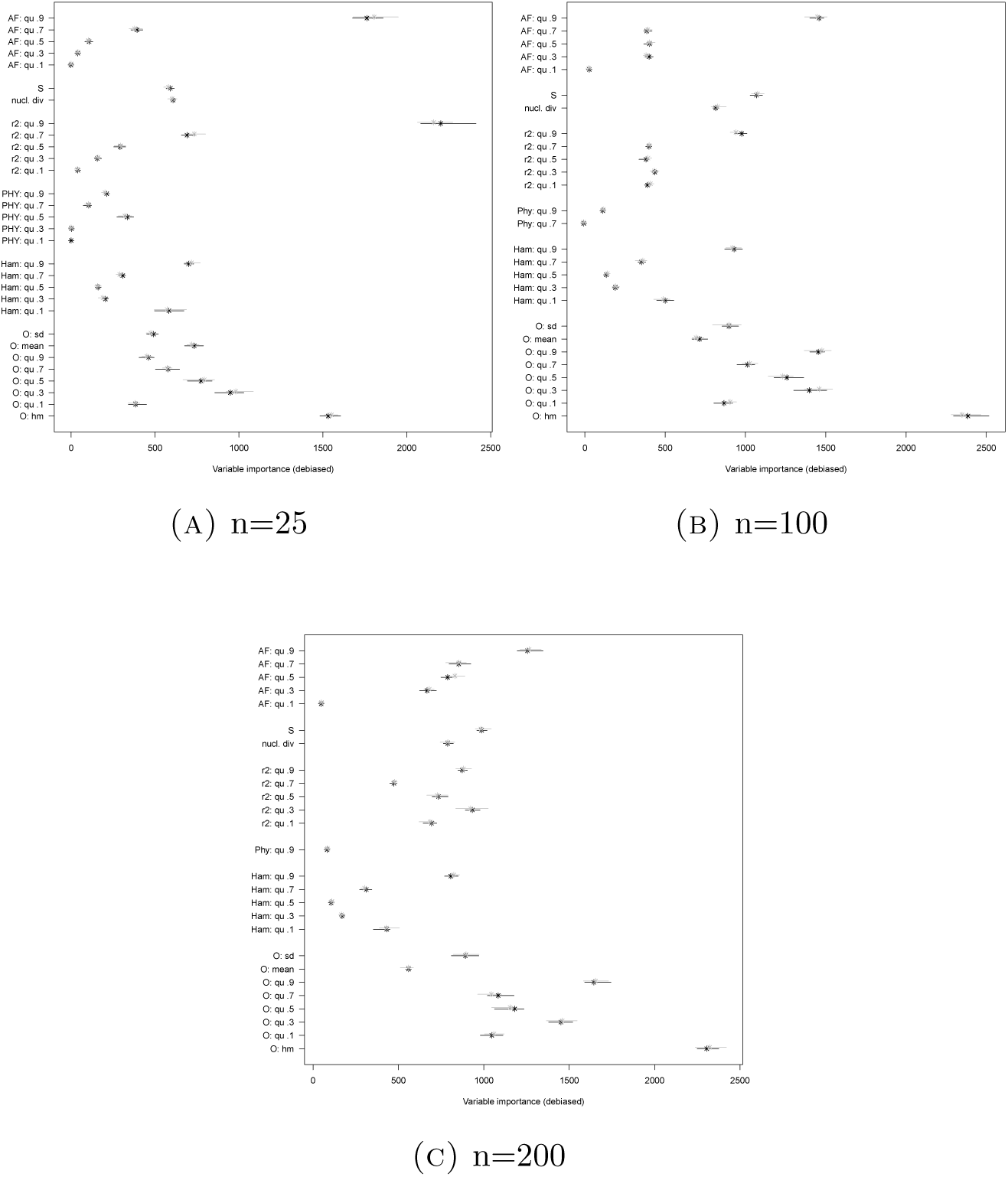
Variable importances for the model comparison between 𝕂-*exp* and 𝔹 (Scenario 2) using **O, Ham, Phy, r**^***2***^ and **AF**+. See Table 7 for the legend.

#### 2.2.2. Adding more quantiles

Above, we followed [KVS^+^17] and used relatively few quantiles of each statistic. Does adding further quantiles decrease errors meaningfully? Especially for **AF**, an alternative is to directly use the full site frequency spectrum **SFS** and/or even further summaries thereof. First, we assessed the effect to use the full site frequency spectrum as well as two further summaries, Tajima’s *D* and Fay and Wu’s *H*, instead of only quantiles of the allele frequencies, see Figure 4.

**Figure 4.**
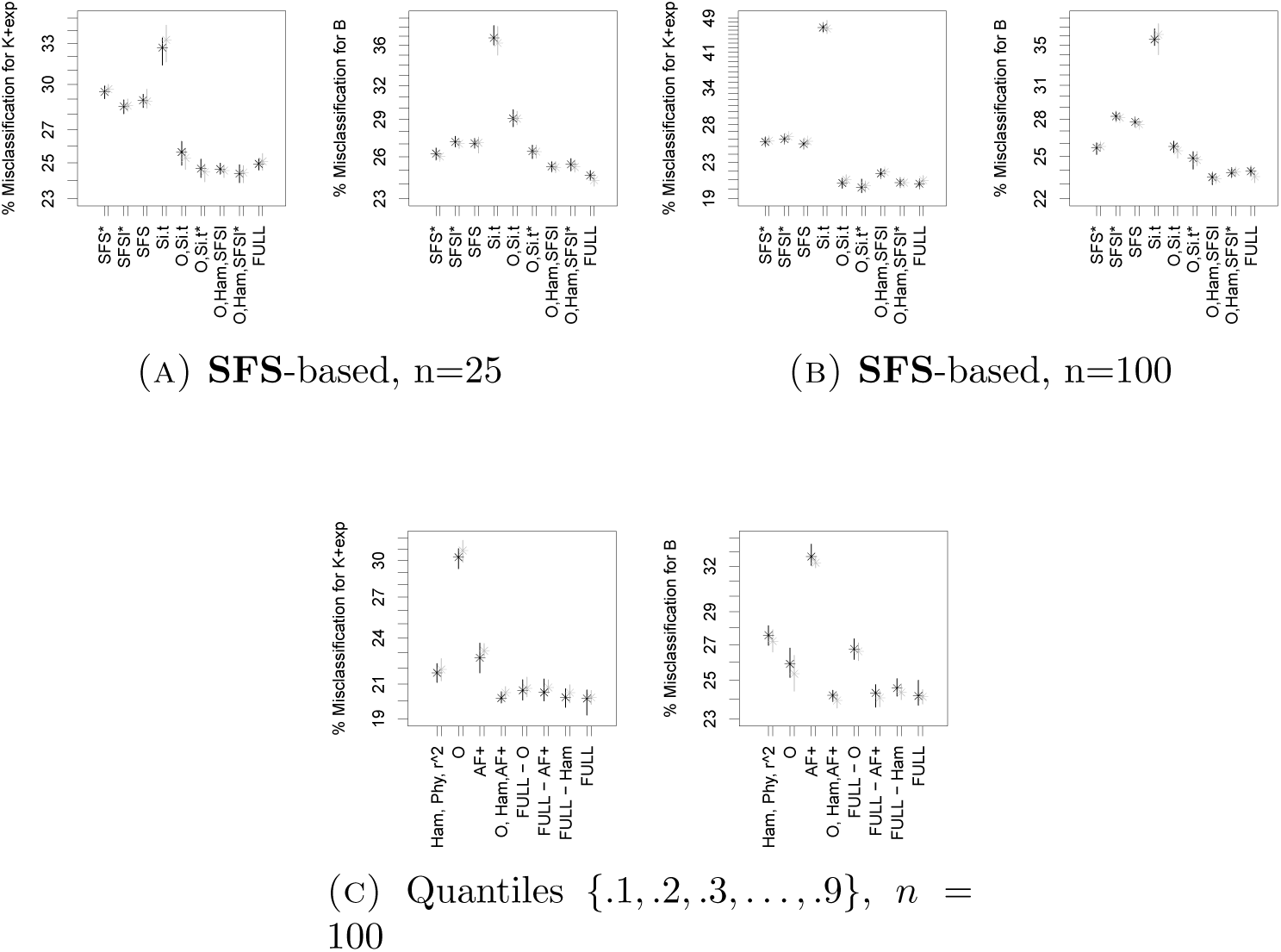
Classification errors (on a log-scale) of model comparison between 𝕂-*exp* and 𝔹 (Scenario 2). See Table 7 for the legend. The full set of statistics consists of (A), (B) **r**^***2***^, **Phy, Ham, O**, *S, π, D, H* and **SFS** and (C) **r**^***2***^, **Phy, Ham, O**, *S, π* and **AF**, but represented by more quantiles. We denote *Si.t* = (*S*_1_, *S*_15+_).

If model selection is performed using only allele frequencies (plus *S* and *π*) or only the site frequency spectrum, the latter decreases errors meaningfully (mean OOB error reduced by ≥2%). The mean error does only depend meaningfully on whether we add *S*\* := (*S, π, D, H*) to the SFS and whether we additionally lump its tail if sample sizes are high enough (mean OOB changes ≤0.2% for *n* = 25, ≥1% for *n* = 100). While adding *S*\* to the SFS reduces errors (mainly due to less misclassification of Beta-*n*-coalescents), lumping the SFS on top of this increases the error beyond using only the SFS (but still well below using **AF**+).

However, the positive effect of using the (lumped) SFS and further summaries vanishes if it is used to perform model selection alongside **O, Ham** (mean OOB error only reduced by ≤0.1% for *n* = 25 and increased by ≈0.1% for *n* = 100). A similar effect can be observed when we add more quantiles to all sets of statistics: A reduction in classification error by adding more quantiles when using **AF**+ alone (mean OOB error decreased by ≈1.2%) as summary statistics vanishes when using **O, Ham, AF**+ as summary statistics and recording more quantiles for all three sets of statistics. This shows that while the (lumped) SFS does contain more useful information to distinguish the different genealogy models than only its coarse representation with 5 quantiles, this information is redundant when the further statistics are included, which is also expressed in low variable importances for most entries of the (lumped) SFS, see Figure 5. From the lumped SFS, *S*_1_ and *S*_15+_ are, especially for *n* = 100, the entries that have the highest importance. These statistics, divided by *S*, constitute the singleton-tail statistic of [Kos18] that is powerful to distinguish between 𝕂-*exp* and 𝔹 for multilocus data. Interestingly, using only these two statistics for model selection fares much worse than using the whole SFS in our one-locus approach, but using them together with **O** or with **O, S**, *π*, **D, H** shows their positive contribution to select models when using the SFS (mean OOB error only increased by 1% resp. 0.3% compared to using all statistics for sample size *n* = 100). Still, this result also shows the further positive effect of adding more information contained in the SFS, e.g. *S*, than just *S*_1_ and *S*_15+_, even when other statistics as **O** are added (also reflected by the comparable or higher variable importances of some of the statistics from *S*\*).

**Figure 5.**
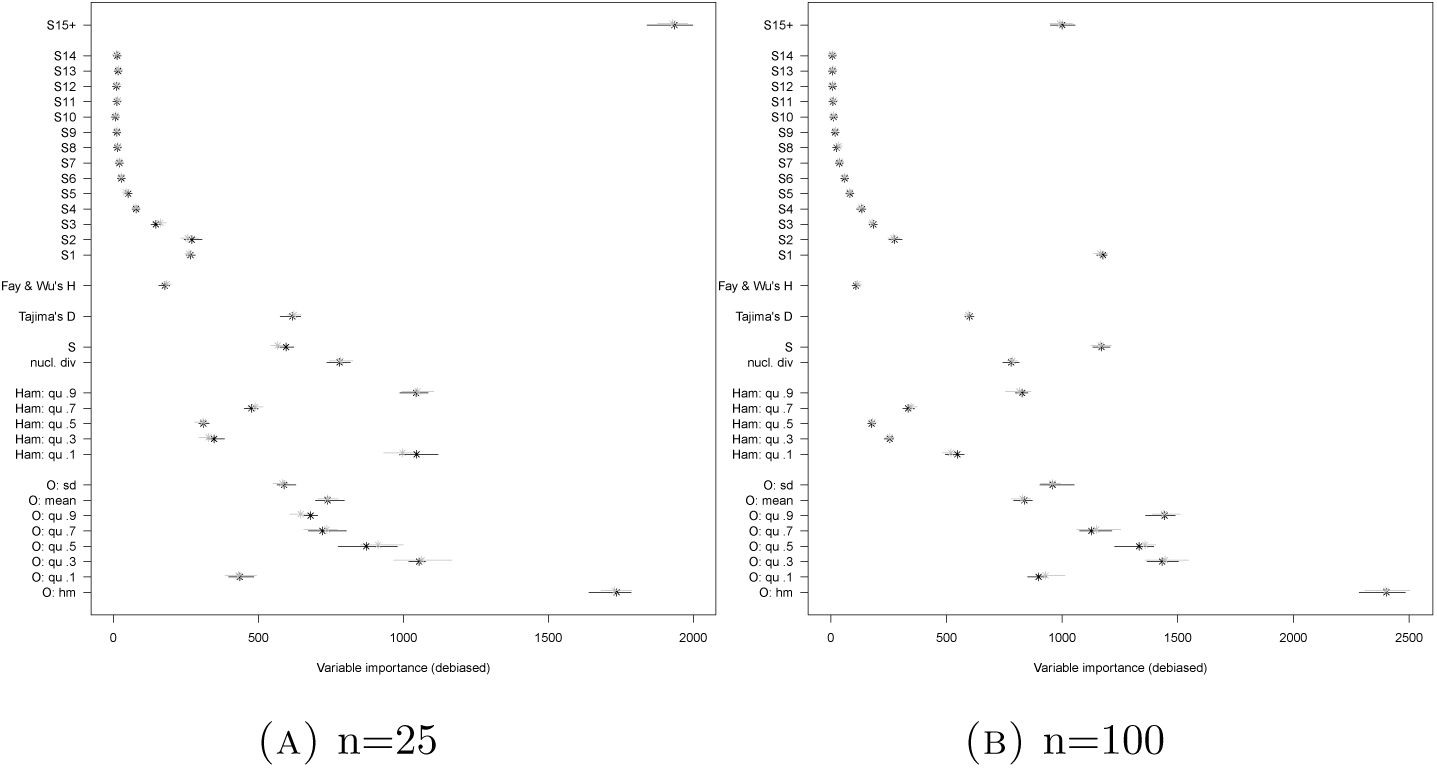
Variable importances for the model comparison between 𝕂-*exp* and 𝔹 (Scenario 2) using **O, Ham, Phy** and and **SFSl**+. See Table 7 for the legend.

An additional insight from this analysis is that **O** is also not a fully reliable set of statistics on its own: using it alone leads to slightly increased error compared to **AF**+ (mean OOB error increase ≈0.4% in the case of increased quantiles for both sets of statistics). However, when not looking at the average error, but the error of misclassifying either 𝕂-*exp* or 𝔹, both sets of statistics act very differently: **AF**+ produces 8% less misclassification of *K*-*exp* than **O**, while **O** decreases misclassication of B by nearly 7%.

#### 2.2.3. Rescaling statistics

The mutation rate strongly affects many of the summary statistics used and thus likely also the model selection, which we also see from the comparison between scenarios 1 and 2. For SFS-based model selection, previous publications scaled the SFS by dividing it by *S* which weakens the effect of mutation rate differences, see e.g. [EBBF15] or the singleton-tail statistics from [Kos18]. Thus, can we also decrease model selection errors by scaling the statistics in our setting? We used scaled versions of the SFS, the Hamming distances and **O** to assess whether using robust inference statistics helps decreasing errors. We compared model inference in Scenario 2 (model classes 𝕂 -*exp* vs. 𝔹) with both the scaled and unscaled statistics, see Figure 6. Being not influenced strongly by mutation rate changes works if mutation rates are high enough, see Figures A1-A4. However, the information lost by scaling increases error rates slightly, see Figure 6 (mean OOB error increase of ≈1.6% for inferring via **nO, nHam, nSFS** instead of their unscaled versions). While inferring model classes using either **nHam** or **nSFSl** does yield error rates that are essentially unchanged by scaling, there is still some information lost when their combination is used (mean OOB error increase of ≈0.7%). The scaling of **O** by the maximum of the minimal observable clade sizes is relatively crude, which is reflected by an elevated error increase for inferring using **O** compared to the other sets of statistics (mean OOB error increase of ≈ 1.1% when using **O** and ≥ 1.6% when added to other sets).

**Figure 6.**
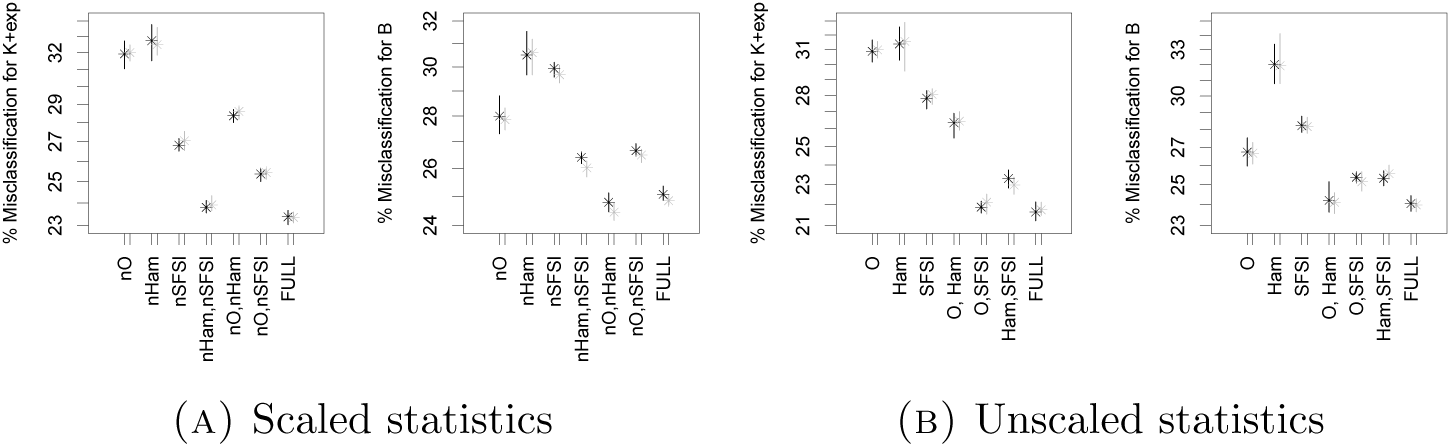
Classification errors (on a log-scale) of model comparison between 𝕂-*exp* and 𝔹 (Scenario 2) with *n* = 100. See Table 7 for the legend. Full set of statistics used: (A) **nO, nHam, nSFSl** (B) **O, Ham, SFSl**.

#### 2.2.4. Low quality statistics

So far, we assumed that one can polarize the data, i.e. that the ancestral and derived allele can be distinguished with certainty. However, this is usually done by comparison with one or more outgroups and is not error-free, see e.g. [KJ18]. Thus, this depends on the availability of suitable outgroups and may not always be possible or precise enough. If mutant and ancestral alleles cannot be distinguished, **O** cannot be computed and, for allele/site frequencies, one can only compute the folded site frequency spectrum **fSFS** (resp. the minor allele frequencies). The other statistics *S, π*, **Phy, Ham** and **r**^***2***^ are still computable. In this case, while error rates are increased compared to when polarized data is available (for mean OOB by > 2% when using **fSFS**+, **r**^***2***^, **Phy, Ham** instead of e.g. **O, Ham, AF**+), there is still a clear benefit of adding statistics to the **fSFS**, see Figure 7(A). Adding **r**^***2***^, **Phy, Ham** to **fSFS**+ reduces model misclassification strongly (≈ 5% mean OOB reduction). Interestingly, adding **Phy, Ham** on top of **r**^***2***^ does help, while again not all entries of the fSFS are well suited to distinguish the model classes. This is also reflected by the variable importances, which are well distributed across the different sets of statistics (except the 0.9 quantile of *r*^2^ having higher importance), while many entries of the fSFS have very low importance, see Figure 8(A). Other factors may also influence whether all statistics can be measured (well) from SNP data. For instance, if the data are not sequenced with high quality (e.g. low coverage data), singleton mutations, i.e. mutations contributing to *S*_1_ (the number of external mutations), may not be highly reliable due to sequencing errors (or overcompensating for sequencing errors). Thus, we assessed what happens if we remove all singleton mutations before computing the statistics and perform inference with **O, Ham, r**^***2***^, **AF**+, see Figure 7(B). The information of the singleton mutations is important for the model selection between 𝕂-*exp* and 𝔹, reflected by a moderate increase in error rate when comparing to results without removal of them (OOB error increase of ≈2% when comparing inferring via **O, Ham, r**^***2***^, **AF**+ without singletons). In this case, including **O** is important, reflected by the highest increase in mean OOB error for **O** when excluding exactly one set of statistics (followed by **r**^***2***^). That the eligibility of **O** for the model selection is not diminished compared to the scenario with singleton mutations is not very suprising, since **O** does not take into account singleton mutations and thus is identical in both scenarios. However, the information available changes by the exclusion of singleton mutations, which strongly affects the ranking of statistics by their importance, see Figure 8(B). Interestingly, the harmonic mean from **O** is not the most important statistic any more (high quantiles of **AF**+, **r**^***2***^ have higher variable importance).

**Figure 7.**
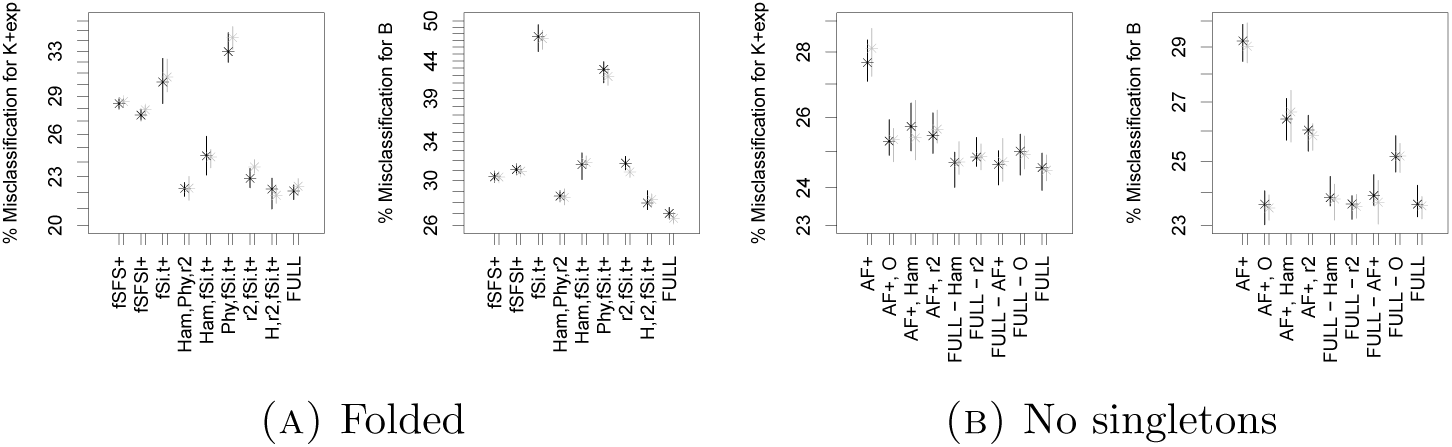
Classification errors (on a log-scale) of model comparison between 𝕂 -*exp* and 𝔹 (Scenario 2) with *n* = 100. See Table 7 for the legend.

**Figure 8.**
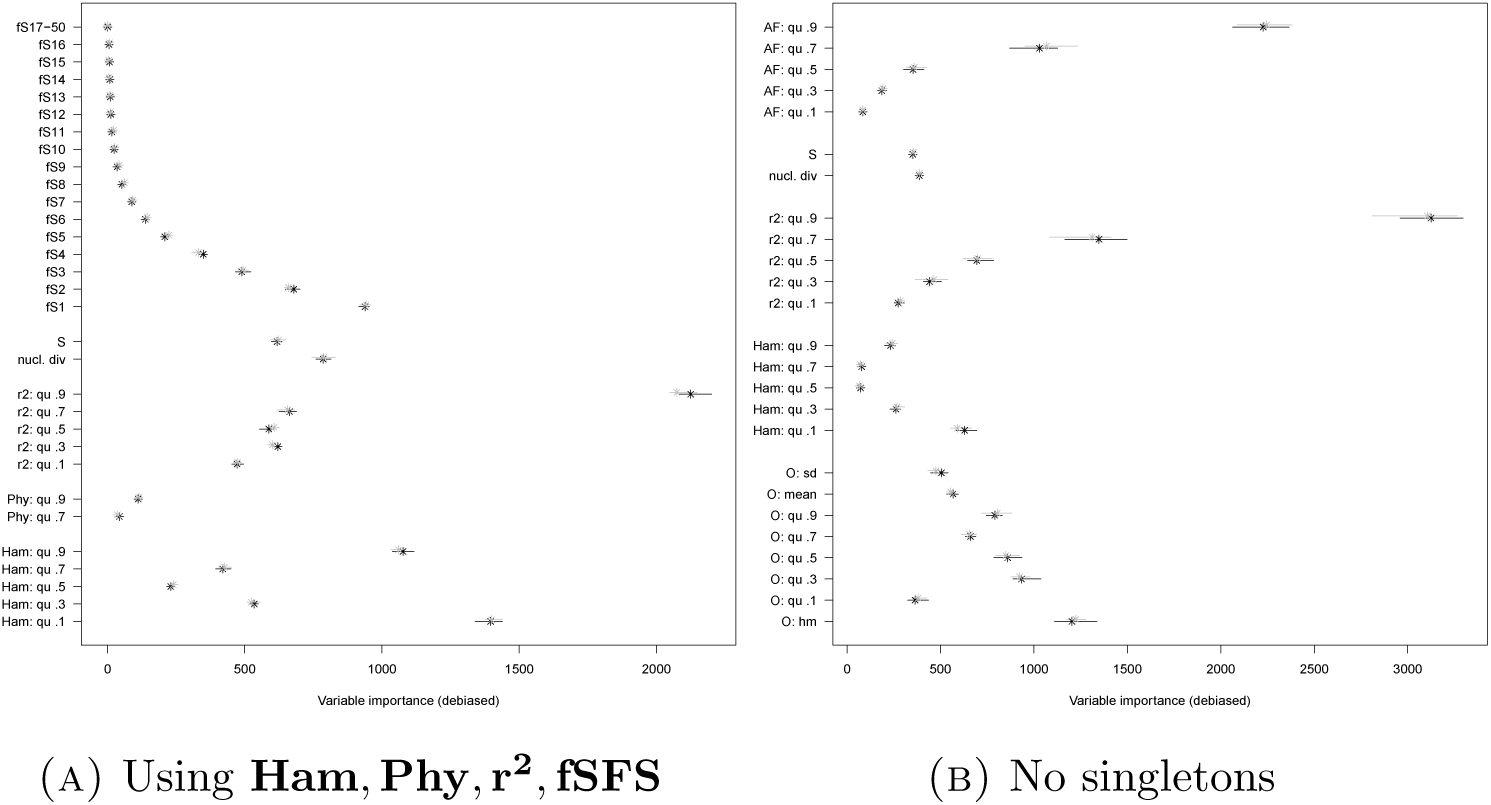
Variable importances for the model comparison between 𝕂-*exp* and 𝔹 (Scenario 2). See Table 7 for the legend. (A) ‘fS17-49’ shows the range (and means) of the importance scores of the variables *f S*17, …, *f S*49 combined.

#### 2.2.5. Scenario 3: adjusting to data

Some bacterial pathogens are haploid species with a single-locus genome, neglectible amount of recombination, and potential for multiple mergers, for instance due to rapid selection or quick emergence of specific strains. Here, we look at a scenario, which we denote by Scenario 3, matching a sample of *Mycobacterium tuberculosis* from [LRP^+^15] regarding sample size and number of mutations. Since the Watterson estimator 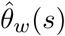 will be not estimating the true mutation rate perfectly, we allow *θ* to fluctuate around the Watterson estimate in the models, see Section 1.2. Misclassification is still common in Scenarios 1 and 2. In Scenario 3 however, classification errors drop considerably, while adding statistics to the SFS strongly decreases misclassification, see Figure 9. As in Scenario 2, **O** is the meaningful addition to **AF**+ to decrease errors and adding further statistics has virtually no effect (adding **O** reduces mean OOB error from ≈ 19% to 9%, other combinations have increased errors).

**Figure 9.**
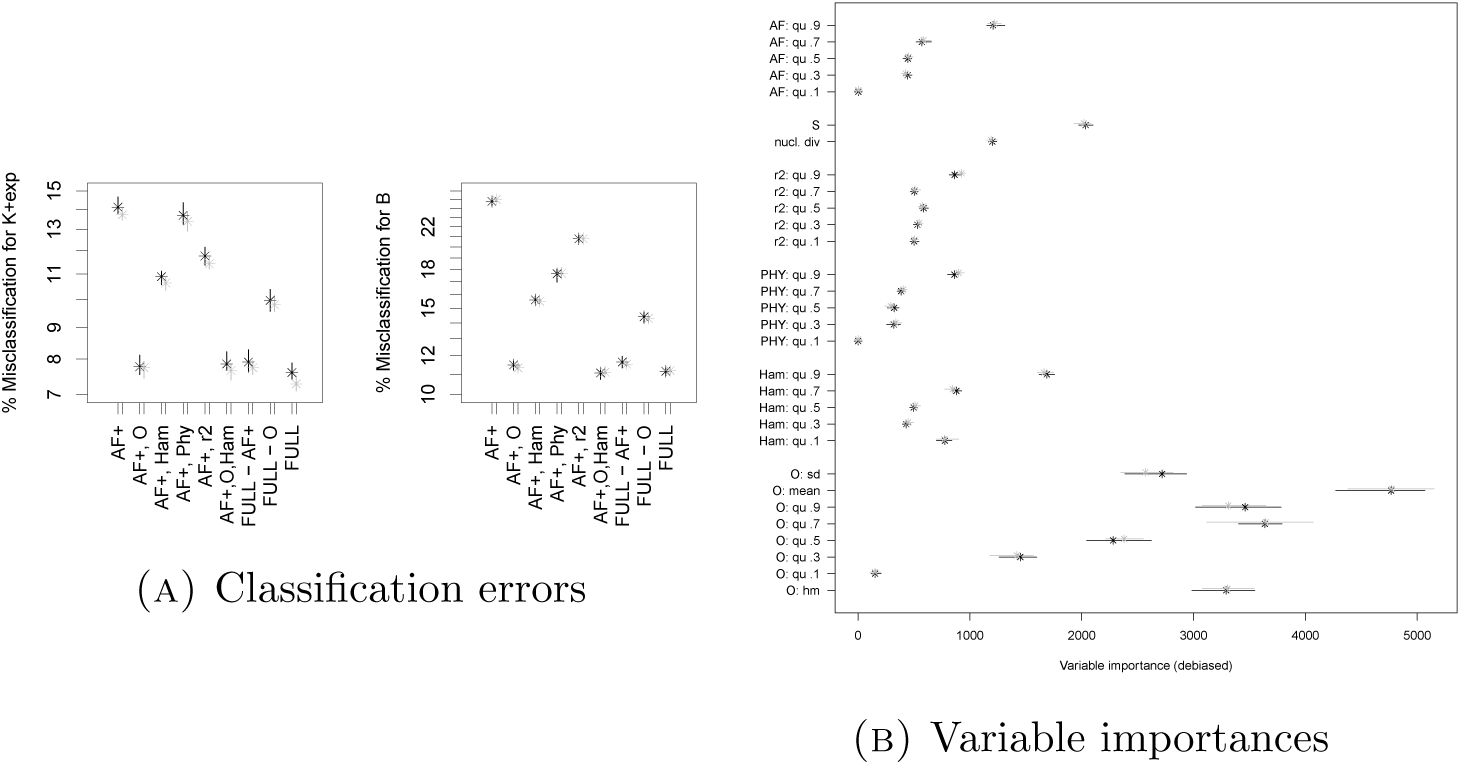
Classification errors (on a log-scale) and variable importances of model comparison between 𝕂 -*exp* and 𝔹 (Scenario 3). See Table 7 for the legend. Full set of statistics **Phy, Ham, O, r**^***2***^ and **AF**+.

### 2.3 Scenario 2: Comparison with Dirac and Beta-Ξ-*n*-coalescents

We consider again Scenario 2 with *n* = 100, where we subsequently add other model classes.

#### 2.3.1. Adding 𝔻 and 𝔻-exp

First, we perform three- and fourfold model selections, the results are shown in Figures 10 and 11. If we compare with the (pairwise) model selection between 𝕂-*exp* and 𝔹 (Figure 2(B)) with the threefold comparison 𝕂-*exp* vs. 𝔹 vs. 𝔻, we see that adding 𝔻 leads to more frequent misidentification of 𝔹, but not of 𝕂-*exp*. The model class 𝔻 is easier to identify than the other two classes, having a far smaller misidentification probability. When we further add 𝔻-*exp*, we see that misidentification for 𝔻 is quite common, while misidentification of 𝕂-*exp* and 𝔹 is essentially unchanged compared to the threefold comparison. This is due to the misidentification of 𝔻-*exp* as 𝔻 and vice versa, see Table A3. For the threefold model comparison, the statistics in **O** have the strongest effect when added to **AF**+, and they cause higher error increase when left out of the full set of statistics (even higher that wheb withdrawing **AF**+). The positive effect of **O** is remaining for the fourfold comparison, but for the classification erros of 𝔻-*exp* which are increased drastically by the addition of any statistic to **AF**+. These findings are also mirrored by the variable importances. The variable with highest importance is again the harmonic mean of the minimal observable clade sizes, and statistics from **Phy, Ham** are not suited well to distinguish the hypotheses (at least if other statistics are available).

**Figure 10.**
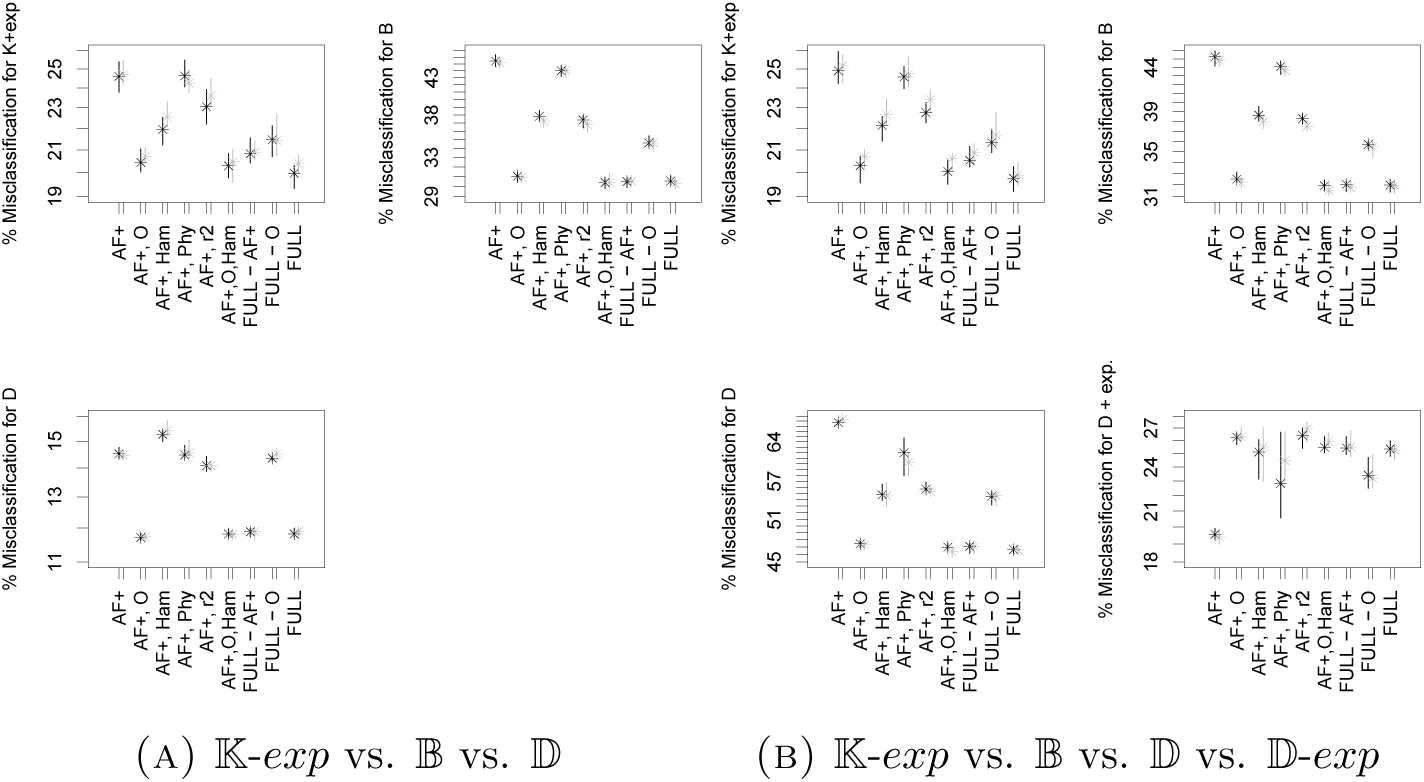
Classification errors (on a log-scale) of model comparisons in Scenario 2 with *n* = 100. See Table 7 for the legend. The full set of statistics consists of **r**^***2***^, **Phy, Ham, O** and **AF**+

**Figure 11.**
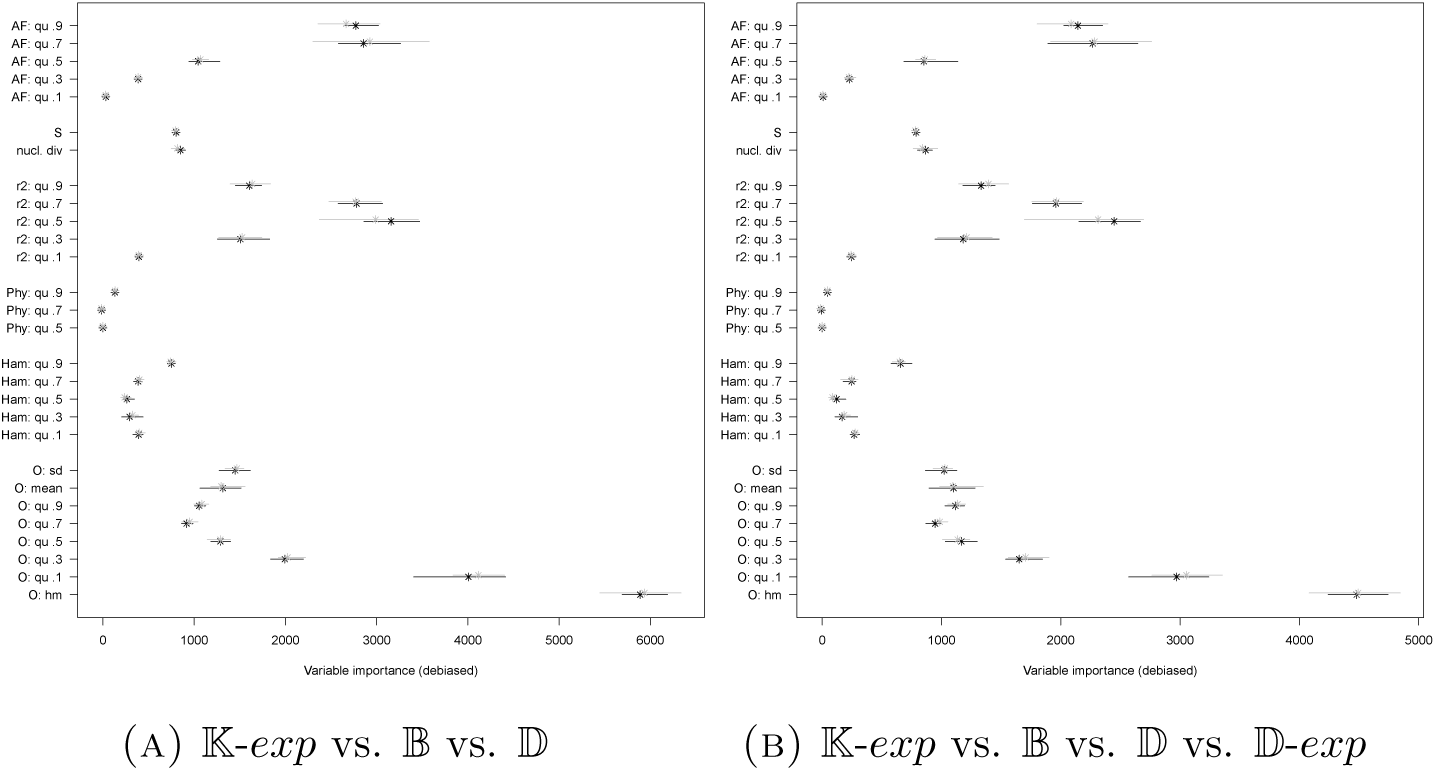
Variable importances in Scenario 2, *n* = 100. See Table 7 for the legend.

How do model selection errors and variable importances change if we consider other pairwise model selections? We additionally performed model selection for 𝕂-*exp* vs. 𝔻 and for 𝔹 vs. 𝔻. The results are shown in Figures 12 and 13. While again **AF**+, **O** (and **Ham**) has essentially the same classification errors as the full set of statistics, for distinguishing the Dirac coalescent **O** is a very suitable addition, both reflected by classification errors and variable importances, where for the latter the harmonic mean from **O** is again the statistic with highest importance. As already known from earlier works, e.g. [EBBF15], the Dirac coalescent is easy to distinguish both from 𝕂-*exp* and 𝔹, reflected by the low misclassification rates compared to the model selection between 𝕂-*exp* and 𝔹.

**Figure 12.**
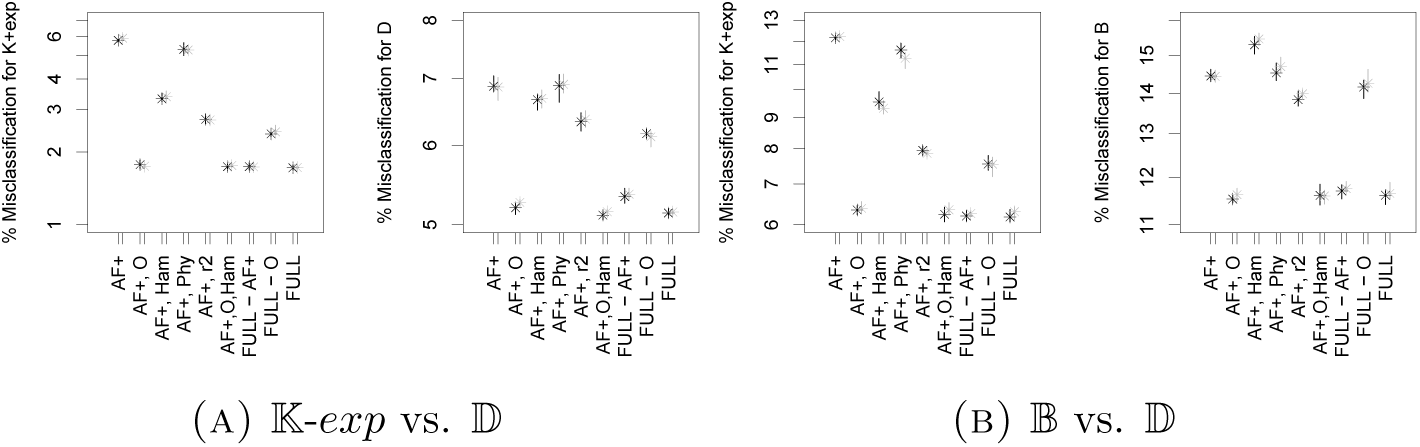
Classification errors (on a log-scale) of model comparisons in Scenario 2 with *n* = 100. See Table 7 for the legend. The full set of statistics consists of **r**^***2***^, **Phy, Ham, O** and **AF**+

**Figure 13.**
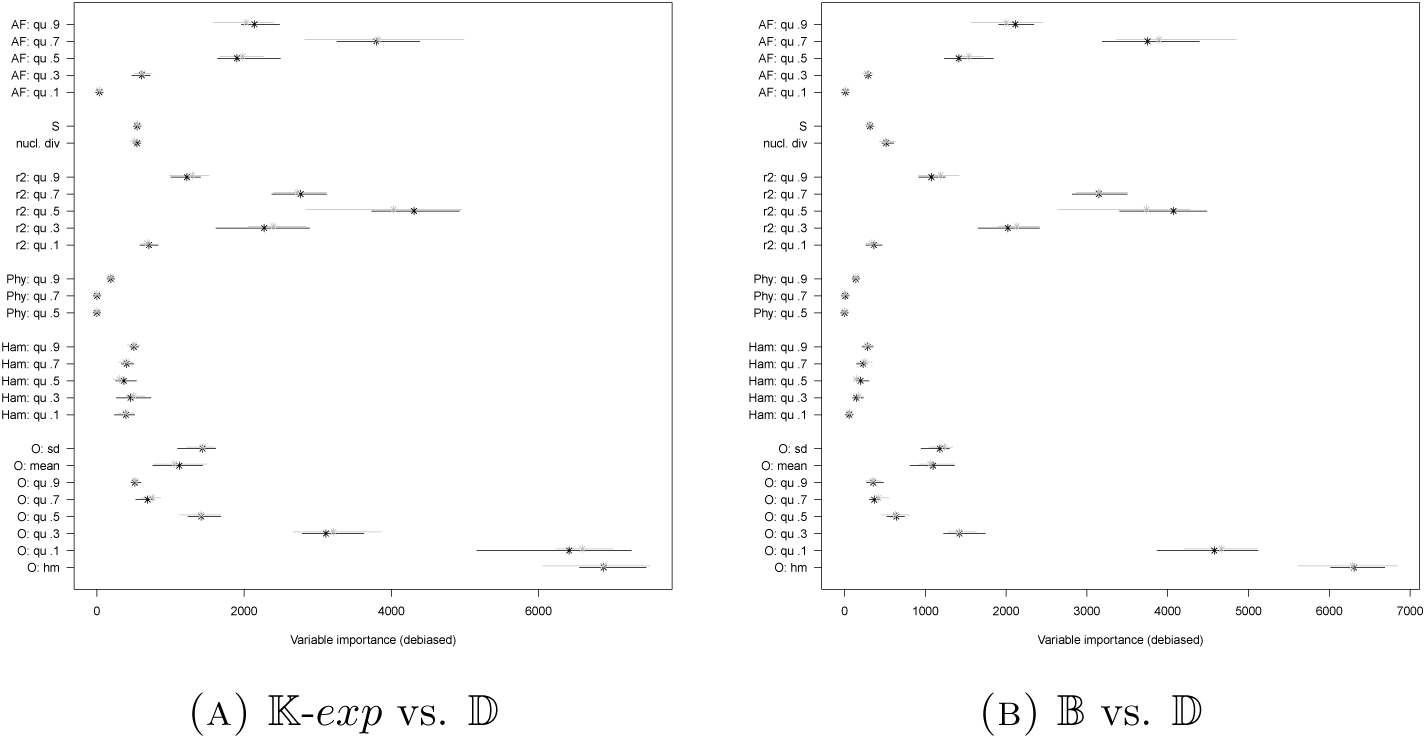
Variable importances for model comparisons in Scenario 2 with *n* = 100. See Table 7 for the legend.

#### 2.3.2. Adding Ξ-𝔹

On the other hand, if we instead add Ξ-𝔹 as a model in Scenario 2, we see a very strong increase in misclassification rates, see Figure 14. The high chance of misclassifying 𝔹 as Ξ-𝔹 stands out, see the results of the pairwise model selection approach in Figure 15.

**Figure 14.**
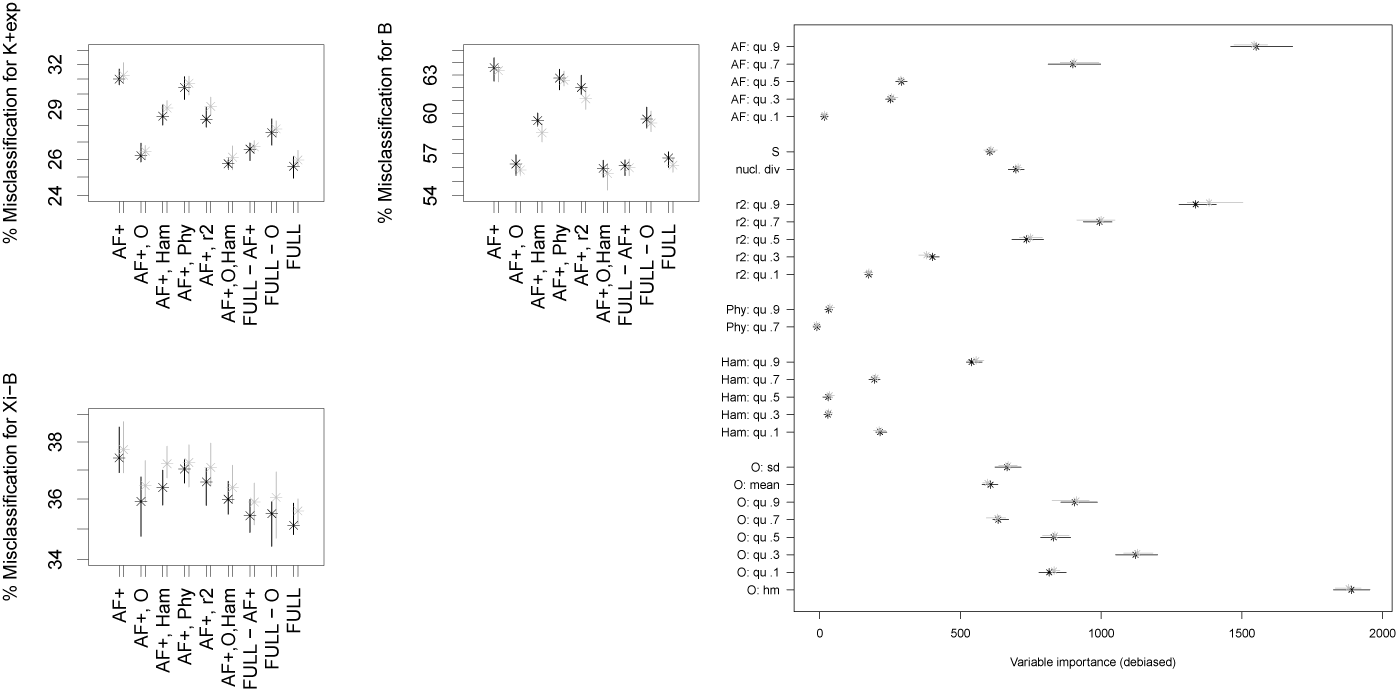
Classification errors (on a log-scale) and variable importances for model comparison 𝕂-*exp* vs. 𝔹 vs. Ξ-𝔹 in Scenario 2 with *n* = 100. See Table 7 for the legend. The full set of statistics consists of **r**^***2***^, **Phy, Ham, O** and **AF**+

**Figure 15.**
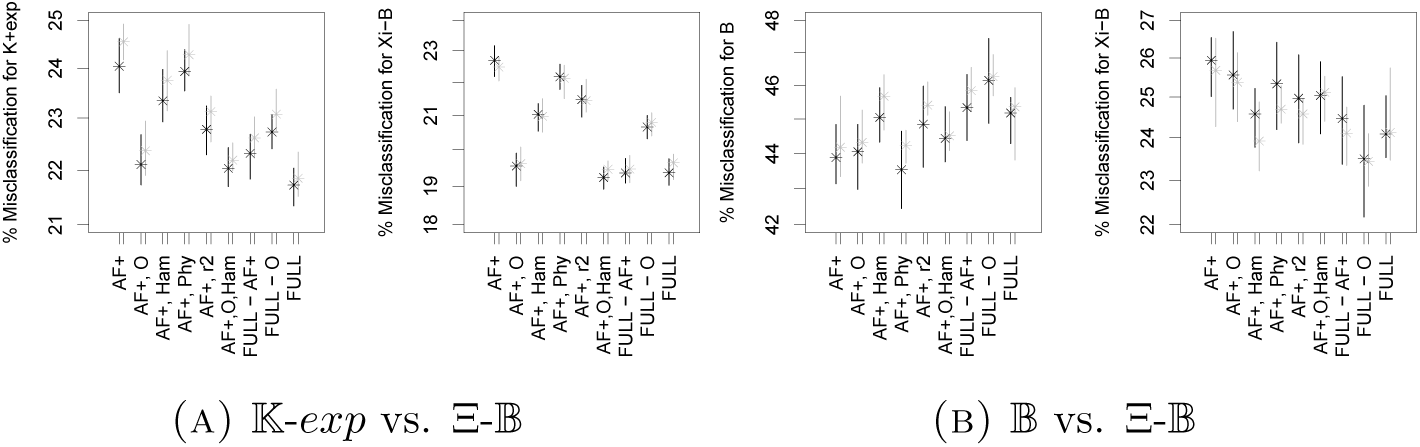
Classification errors (on a log-scale) of model comparisons in Scenario 2 with *n* = 100. See Table 7 for the legend. The full set of statistics consists of **r**^***2***^, **Phy, Ham, O** and **AF**+

Again, performing the model comparison based on **O** and **AF**+ does not increase misclassification meaningfully compared to using all statistics (if one does not simply choose not to compare models due to very high errors). While for the threefold comparison, the most important statistic still is the harmonic mean from **O**, followed by statistics from **AF, r**^***2***^, **O**, this changes if one looks at the pairwise comparisons. There, **O** only plays an important, yet diminished role for distinguishing Ξ-𝔹 from 𝕂-*exp*, it seems to play much less of a role for distinguishing Ξ-𝔹 from 𝔹 (Figure 16). This is not entirely mirrored by the error rate changes between different sets of summary statistics, see Figure 15, but the differences between most sets of statistics for these scenarios are small to begin with.

**Figure 16.**
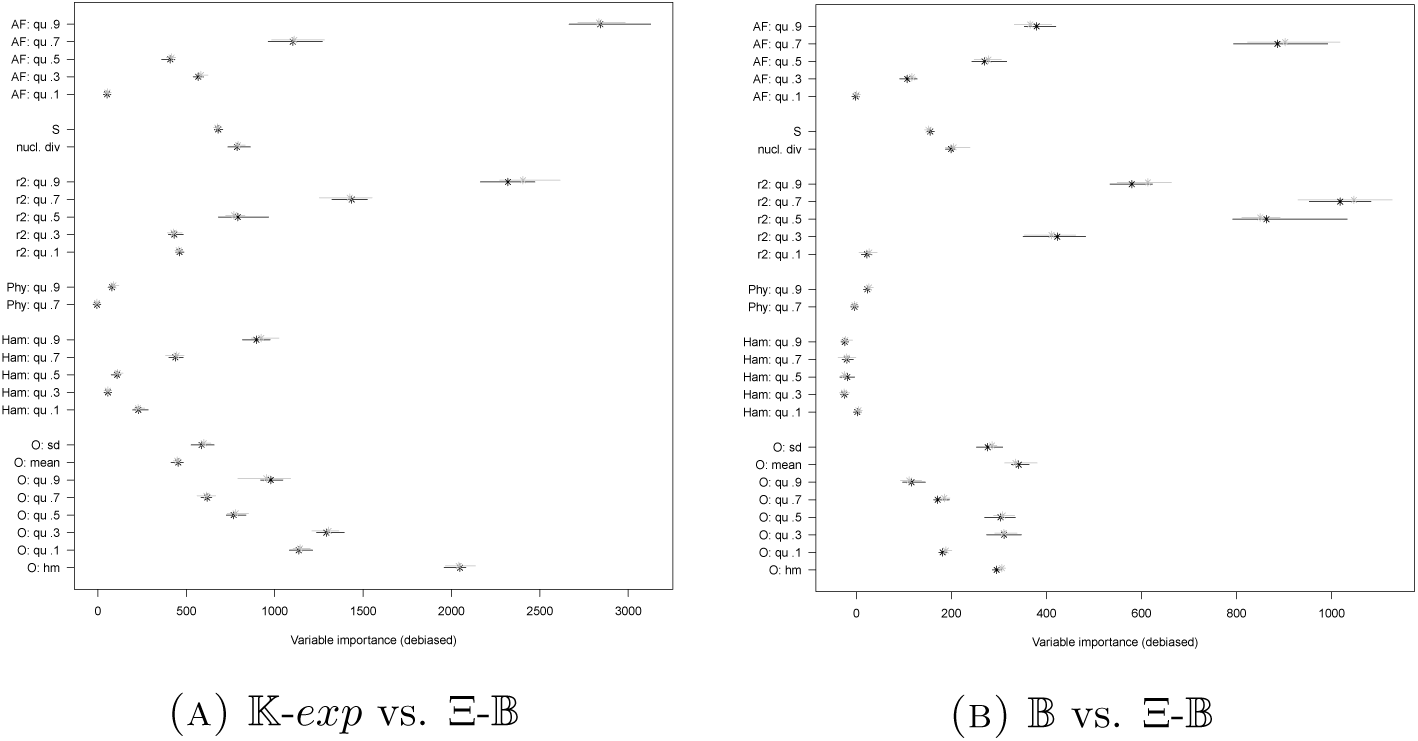
Variable importances for model selection in Scenario 2, *n* = 100. See Table 7 for the legend.

#### 2.3.3. Misspecification of coalescent models

Our results imply that the distribution of diversity statistics within different (parametric) subclasses of Ξ-*n*-coalescents can both be rather similar or very different. This also raises the question whether misspecifying the exact multiple merger model as a different multiple merger model increases model selection errors. We analyze this by allocating simulations from one Ξ-*n*-coalescent class to either 𝕂-*exp* or another Ξ-*n*-coalescent, see Section A.1 for details. For instance, we build a random forest from simulations of 𝕂-*exp* and 𝔹, but sort simulations from 𝔻 into these two classes via ABC. The results are shown in Table 8. They highlight that multiple merger *n*-coalescents do not generally differ in the same way from 𝕂-*exp*. While 𝔻 gets nearly always identified as 𝔹 instead of 𝕂-*exp*, this is not symmetric: 𝔹 gets considerably more often identified as 𝕂-*exp*. Unsurprisingly, given the high classification errors between 𝔹 and Ξ-𝔹, both models get more often identified as the other multiple merger model, with chance around 0.7, in contrast to as being identified as 𝕂-*exp*. Still, model misspecification comes with a cost: if we use 𝔹 as a proxy model for Ξ-𝔹 and have to allocate a true realization of Ξ-𝔹 to either 𝔹 or 𝕂-*exp*, this leads to considerably higher classification errors (though they should still be manageable) than when performing the model selection with the correct model Ξ-𝔹, see Figure 15. However, in this case only considering simulations that can be allocated to a model class with high posterior probability lowers errors, but at the (different) cost of being unable to allocate a high proportion of observed data. To a degree, one can try to balance cost and benefit here, e.g. retaining only simulations from Ξ-𝔹 with posterior > 0.64 for either 𝔹 or 𝕂-*exp*, which is fulfilled by a proportion of 0.72 of simulations, leading to a decreased identification chance as 𝕂-*exp* of 0.18, which is comparable to the misidentification rate in the model selection using the correct model Ξ-𝔹.

**Table 8.**
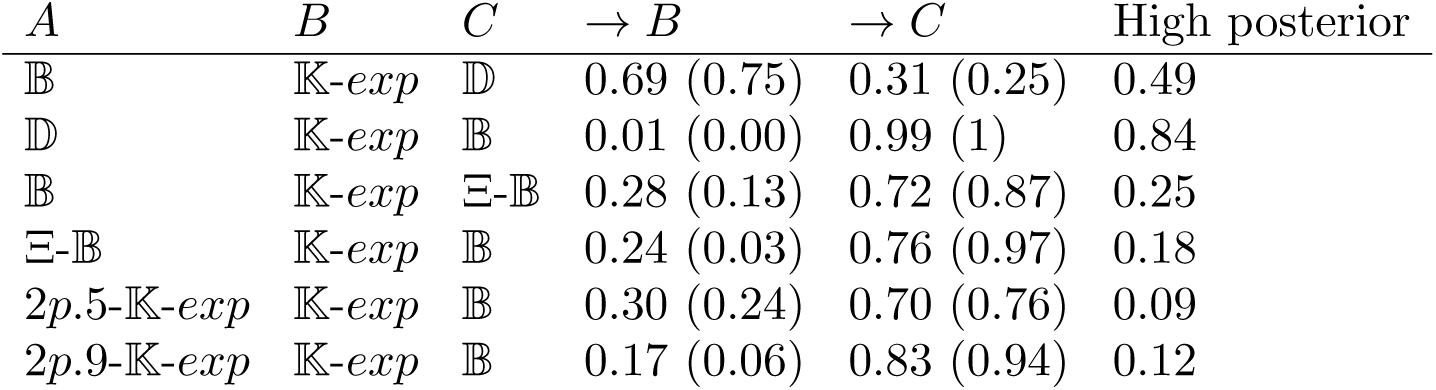
*→B, →C*: Proportion of simulations from model class *A* allocated to two other model classes *B* and *C* in Scenarios 2 and 4. In parentheses: same proportion only among high posterior allocations. High posterior allocations are those with posterior probability > 0.9, the column ‘high posterior’ shows their proportion among all simulations from *A*. Random forests are based on **r**^***2***^, **Phy, Ham, O, AF**+. Results are rounded to two digits.

### 2.4. Scenario 4: The influence of population structure

For two very simplistic scenarios with two subpopulations and low symmetric migration rates, where either equal samples or skewed samples are taken from the subpopulations, we find that model classes 𝕂-*exp*, 𝔹 and 2*p*.5/2*p*.9-𝕂-*exp* with population structure can be distinguished with a slightly higher error than 𝕂-*exp*, 𝔹 and 𝔻, but better than 𝕂-*exp*, 𝔹 and Ξ-𝔹 (averaged over model classes), see Figure 17. As statistics, we use **AF**+, **Ham, Phy, r**^***2***^, **O**. Again, for K-*exp* and B, we see the highest error reduction when coupling **O, AF**+, **Ham**, essentially with the same error rates as when using all statistics. For the structured population models, the highest error reduction is obtained using **Ham, Phy, r**^***2***^, **O**. Dismissing **O** increases the mean OOB error rate from ≈1% (when considering 2*p*.9-K-*exp*) to ≈1.7% (when considering 2*p*.5-𝕂-*exp*). Due to their known potential to distinguish demography from population structure ability, we also checked whether performing model selection using Tajima’s *D* and Fay and Wu’s *H* together with the SFS, *π, S*, instead of **AF**+ lowers misclassification,, see Figure A9. For both population structure choices, combining either **AF**+ or {**SFS**, *π, S, D, H*} with **Ham, Phy, O, r**^***2***^ essentially produces identical errors (using **SFS** decreases mean OOB error further by. ≤1%). The same holds true when using **O, Si.t***\** or **O, AF**+ (the latter decreases mean OOB error by ≤ 1%).

**Figure 17.**
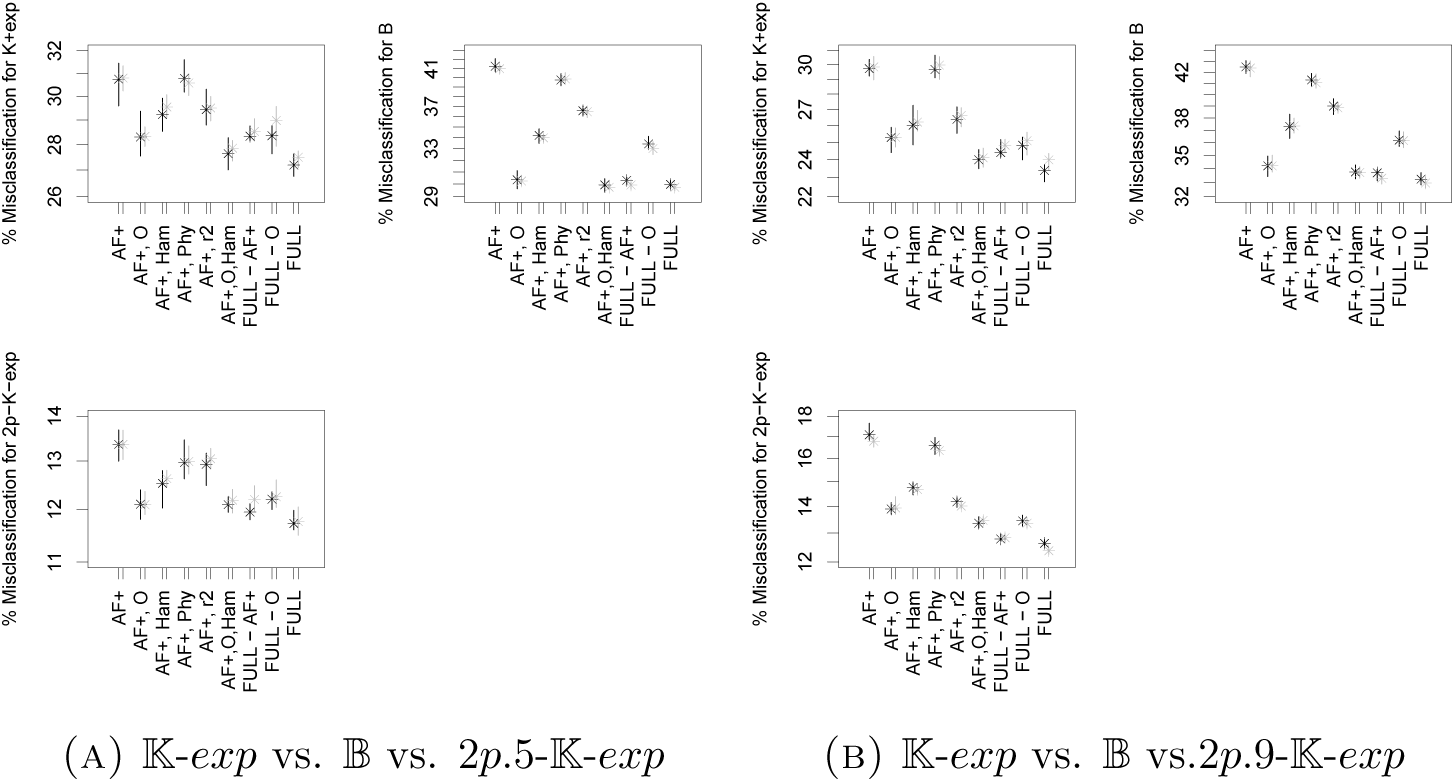
Classification errors (on a log-scale) in Scenario 4, *n* = 100. See Table 7 for the legend. Full set of statistics includes **r**^***2***^, **Phy, AF**+, **O** and **Ham**.

When looking at variable importances, we see that the importance of the harmonic mean of the minimal observable clade sizes is reduced compared to other scenarios, while high allele frequency and Hamming distance quantiles are elevated, see Figure 18. This may reflect that strong population structure tends to produce long branches in the genealogy connecting the root to the samples from different subpopulations, causing bumps in the allele frequency distribution and an abundance of high Hamming distances.

**Figure 18.**
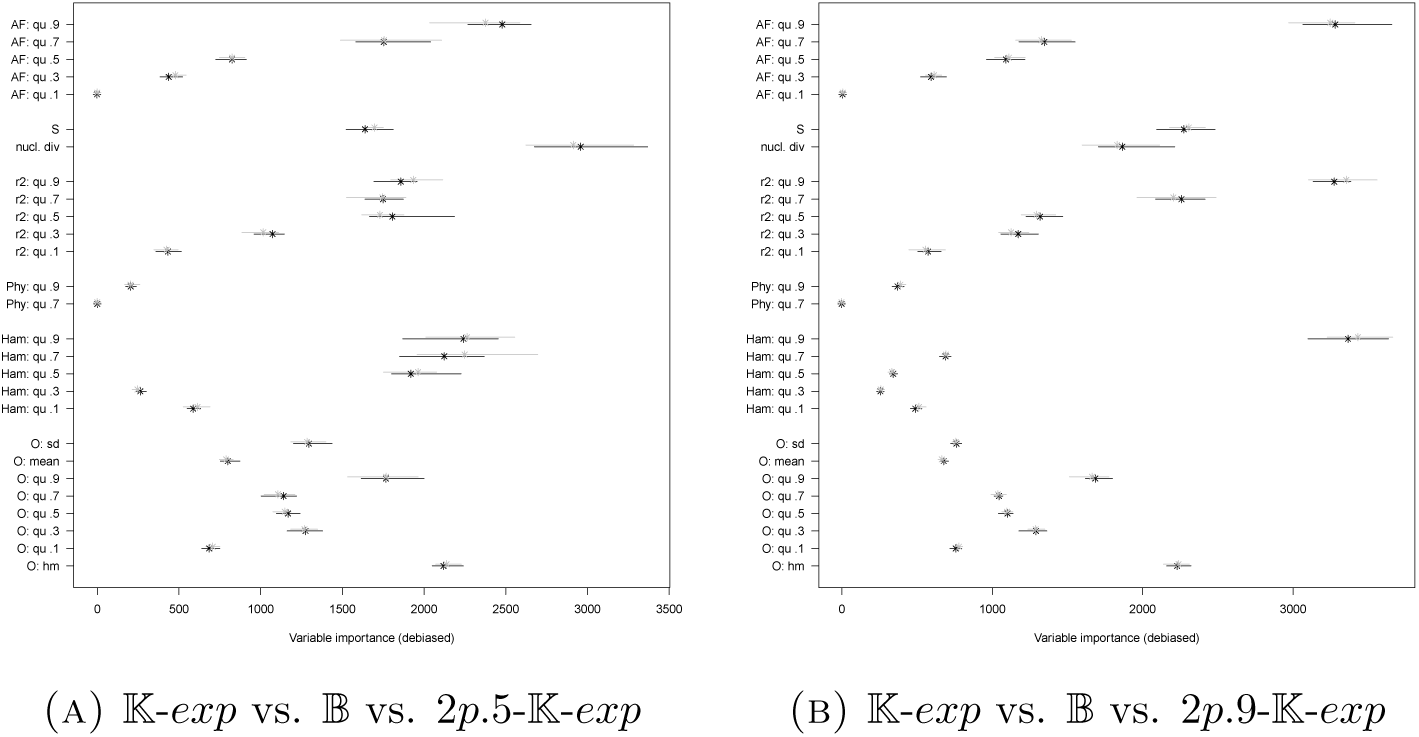
Variable importances in Scenario 4, *n* = 100. See Table 7 for the legend.

The error rates strongly drop for pairwise comparisons between model classes, see Figures 19 and A10. First, have a look at selecting between 𝕂-*exp* with and without population structure. While for equal-sized samples from the subpopulations, adding statistics to **AF**+ does not considerably lower misidentification (< 1% mean OOB error decrease for any addition), this can be the case for unbalanced sampling. For subsample sizes 9:1, we see a drop-off in mean OOB error of ≈3%. In both cases, there is not a strong difference which set of statistics is added. For single variables, importance scores are highest for statistics from **AF**+ and **Ham**, see Figures 20(A) and A11(A). The high importance of statistics from **Ham** is in contrast to most other scenarios (**Ham** shows relatively high importance for *n* = 25 in Scenarios 1 and 2, the latter for 𝕂+*exp* vs. 𝔹). For both equal and unequal sampling from the sub-populations, the .9-quantile statistics from the two sets are among the most important statistics for distinguishing 𝕂-*exp* with and without population structure.

**Figure 19.**
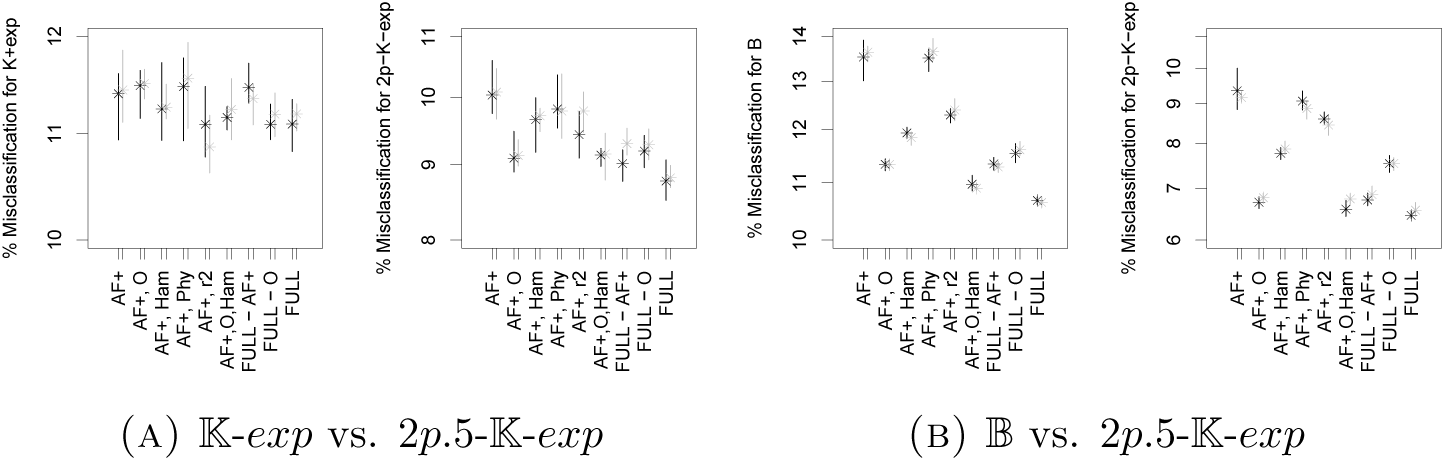
Classification errors (on a log-scale) for model selection in Scenario 4, *n* = 100. See Table 7 for the legend. Full set of statistics includes **r**^***2***^, **Phy, AF**+, **O** and **Ham**.

**Figure 20.**
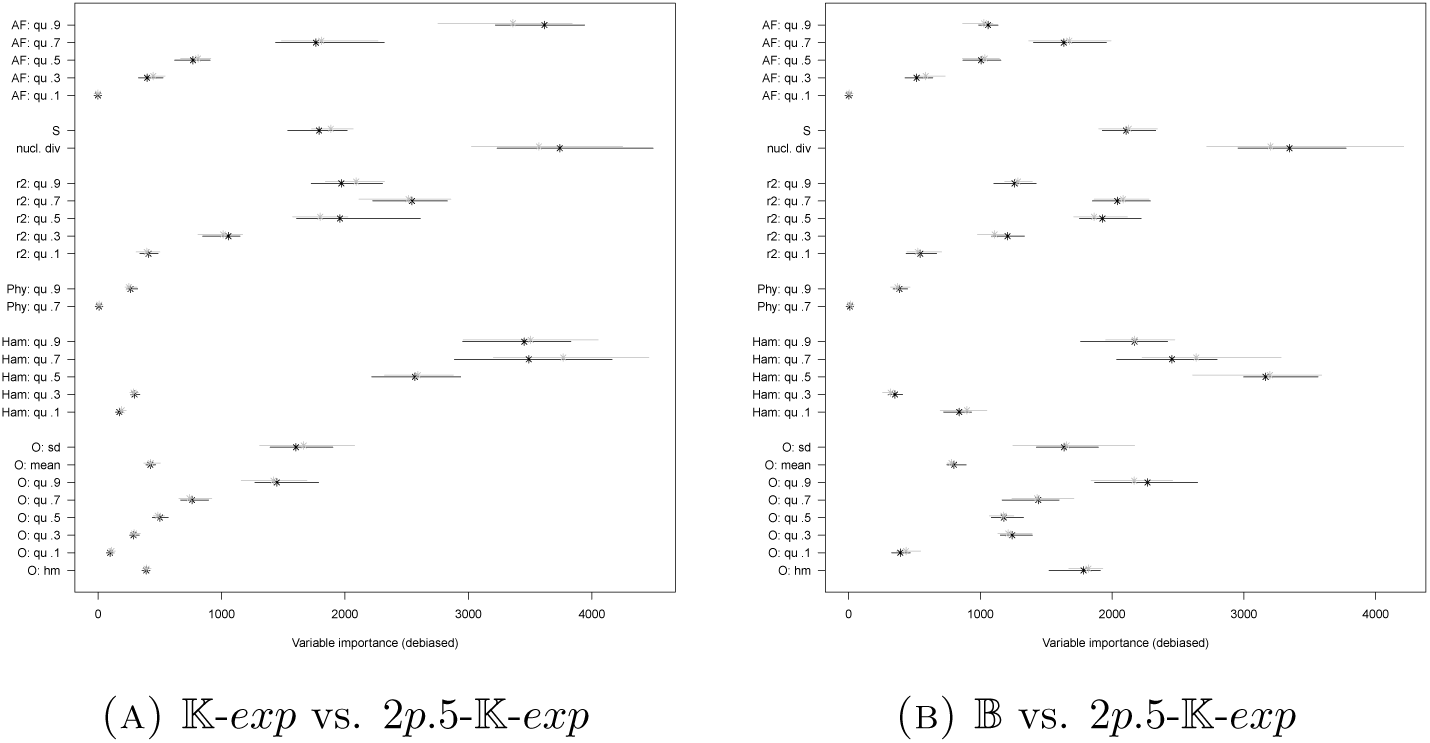
Variable importances in Scenario 4, *n* = 100. See Table 7 for the legend.

For selecting between 𝕂-*exp* with population structure and 𝔹, the additional error decrease when adding statistics to **AF**+ is slightly stronger (≈3% decrease of mean OOB error), which is due to the addition of **O, Ham**, with a stronger effect of the former. For equal-sized supsamples, all statistics but **Phy** have at least some member with reasonable importance (Figure 20(B)), again *π* and (quantiles of the) Hamming distances are among the variables with highest importance. For a sampling ratio of 9:1 from the two subpopulations, the .9 quantiles of Hamming distances and **r**^***2***^ show high importance, as does *S*, see Figure A11(B). Surprisingly, given the stronger effect of **O** on misclassification than the other statistics when added to **AF**+, no single statistic of **O** shows high importance.

While the simple scenario of strong population structure and two subpopulations can be quite well distinguished from 𝕂-*exp* and 𝔹, this only works when population structure is known *a priori*. Indeed, if we allocate simulations from 2*p*.5-𝕂-*exp* to 𝕂-*exp* or 𝔹, i.e. perform model selection between the latter two models based on data that truly follows 2*p*.5-𝕂-*exp*, Table 8 reveals that simulations with population structure tend to be identified as being generated by Beta-*n*-coalescents, especially if sampling is skewed towards one subpopulation. However, to be allowed to ignore population structure for inference, such simulations should be sorted into 𝕂-*exp* with high probability, which is clearly not the case. This effect gets even stronger if one only regards simulations that are allocated to one model class with high posterior probability.

### 2.5. Scenarios 2 and 5: The influence of exponential growth

We further consider Scenario 5, focusing only on comparisons between scenarios with growth and without, see Figure 21. Even for 𝕂 vs. 𝕂-*exp* and high mutation counts, the model with growth is relatively often misidentified, which partly comes from our focus on low to medium growth rates, compared e.g. to [EBBF15]. For instance, subsetting for only growth rates above 20 (and drawing the same number of simulations from 𝕂) leads to strongly reduced classification errors (3% for 𝕂 and 10.3 % for 𝕂+*exp*, single ABC run with both simulation sets combined). Additionally, the larger error for identifying 𝕂-*exp* is also affected by the prior: we have only a relatively small number of simulations from models close to 𝕂 (i.e. with very small growth rates) which are very hard to distinguish from 𝕂. Such simulations, if left out of a bootstrap sample, then get easily sorted into a leaf associated to the more numerous simulations from 𝕂. Generally, for distinguishing these two model classes, adding statistics to **AF** decreases misclassification in the random forest-based ABC model selection only slightly (at most by 1%) and there is no specific set of statistics that has information which is non-redundant for all other sets.

**Figure 21.**
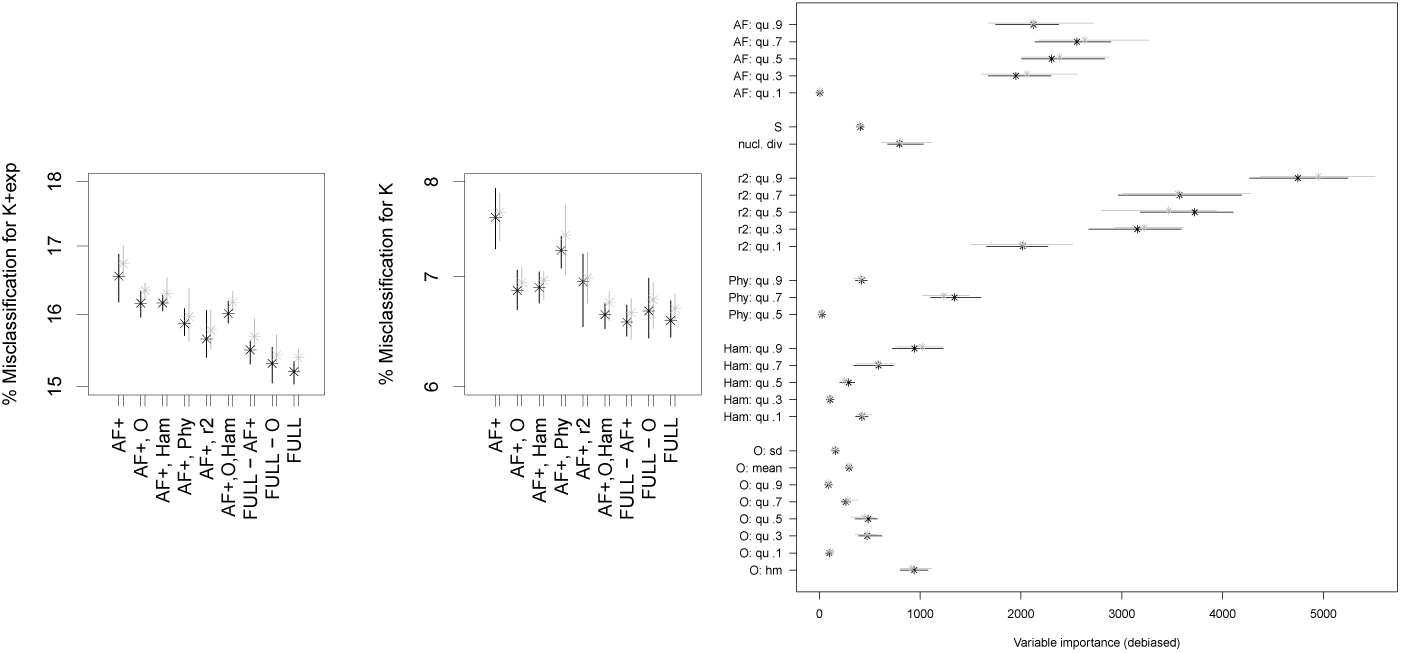
Classification errors (on a log-scale) and variable importances for model comparison 𝕂 vs. 𝕂-*exp* in Scenario 5 (*n* = 100). See Table 7 for the legend. The full set of statistics consists of **r**^***2***^, **Phy, Ham, O** and **AF**+

If one looks at distinguishing 𝔻 from 𝔻-*exp*, now again with the lower mutation rates of Scenario 2, we see increased error probabilities (Figure 22). While 𝔻 could be well distinguished from 𝔹 and 𝕂-*exp*, see Figure 12, it is much more similar in its summary statistics to 𝔻-*exp*. In contrast to model selection between 𝕂-*exp* and 𝕂, adding statistics to **AF** does strongly decrease misclassification for 𝔻, which is directly attributable to adding **O**, but this strategy increases misclassification for 𝔻-*exp*.

**Figure 22.**
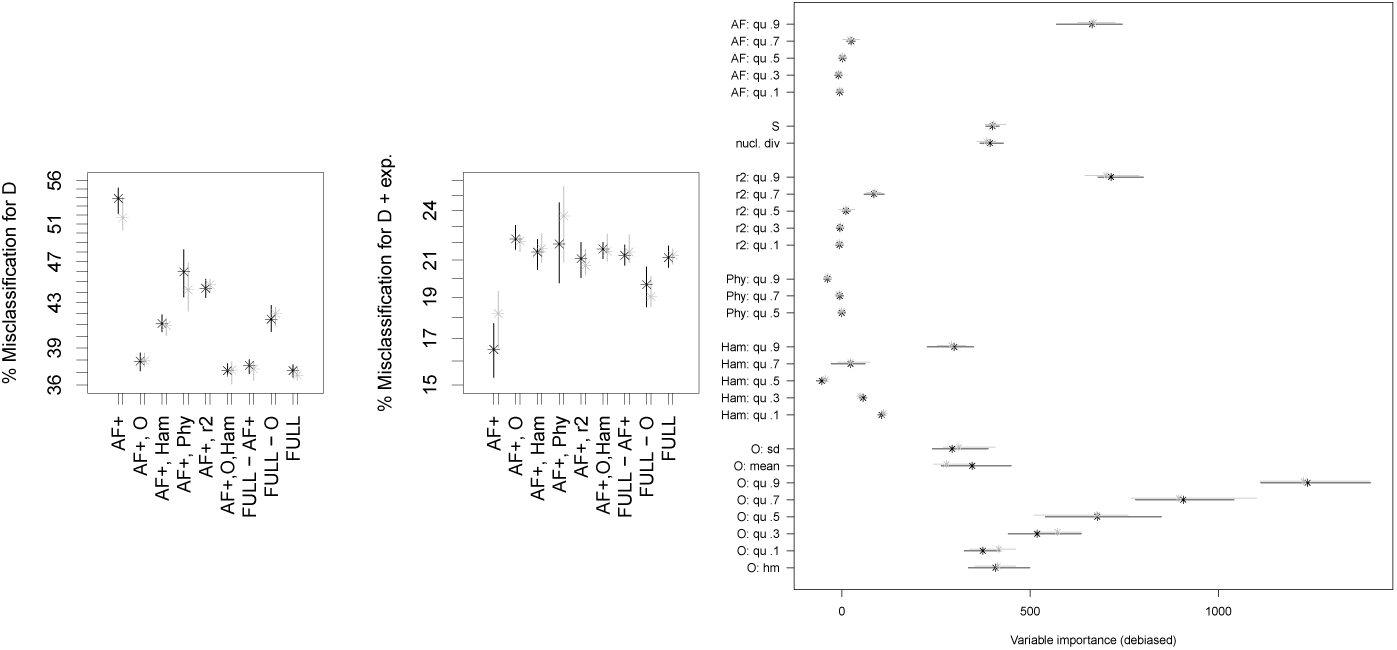
Classification errors (on a log-scale) and variable importances of model comparison 𝔻 vs. 𝔻-*exp* in Scenario 2 for *n* = 100. See Table 7 for the legend. The full set of statistics consists of **r**^***2***^, **Phy, Ham, O** and **AF**+.

For Scenario 5 (Kingman models), **r**^***2***^ (especially the .9-quantile) shows high importance, while for the Dirac models, **O** (especially the .9-quantile) shows high importance (Figures 21 and 22). This is in accordance with the misclassification rates.

## 3. Discussion

Our results highlight that the information not contained in the site frequency spectrum is still highly valuable for distinguishing between different classes of *n*-coalescents, e.g. between exponential growth and multiple merger coalescents. This beneficial effect of using site frequency spectra with other statistics of genetic diversity has been also recently reported for other coalescent models (also incorporating recombination) in [JBA19]. In addition to scenarios with fixed mutation rate ranges across models as in [KVS^+^17], this also holds true for scenarios where mutation rates are chosen so that, on average, each model produces the same number of mutations. While we concentrated on full polarized genetic data, there is still a strong beneficial effect of using more statistics than allele frequencies spectra when data cannot be polarized or if one ignores singleton mutations (due to their susceptibility to being mixed with sequencing errors). The benefit of including statistics not based on the SFS is especially relevant for species with small genomes and few unlinked loci, e.g. for some bacteria, but also likely for some selfing plants and clonally propagating fungi. Giving high enough mutation rates and reasonable sample sizes, as in our Scenario 3, one can distinguish exponential growth from multiple mergers with ease. We emphasize that seeing hundreds of mutations at a single locus in a species with low recombination rates is not unrealistic (though clearly not always the case), we based our setting on real data sets in *Mycobacterium tuberculosis*. While model misidentification becomes less of an issue if the genome consists of many unlinked loci, see [Kos18], this may still be relevant if further evolutionary processes are acting, as discussed in [KWB19]. On the other hand, our results also highlight that some model selections, e.g. between Beta and Ξ-Beta-*n*-coalescents or between Dirac coalescents with and without growth are very noisy with only data from a single locus and likely their selection cannot be trusted.

While adding statistics to the allele frequency spectrum helps with distinguishing genealogy models, the information that helps is already contained in rather few quantiles. Additionally, our analysis shows that the information of the combination of the different statistics allows one to cope with not knowing the mutation rate precisely and that, when statistics are combined, scaling statistics to be non-sensitive to changes in the mutation rate is not necessary (while it still is reasonable when only using allele frequency spectra). Compared with the results in [KVS^+^17], our results show highly increased error rates. This is likely not a consequence of our ABC approach, but of our focus on distinguishing Beta(2 −*α,α*)-*n*-coalescents for *α ∈* [1, 2) from Kingman’s *n*-coalescent with growth, while [KVS^+^17] also add further Beta coalescents that are easier to distinguish from the bifurcating genealogies. Nevertheless, this specific set of Beta(2 −*α,α*)-*n*-coalescents is chosen here because it arises as limit genealogies in an explicit evolutionary model, see [Sch03].

Naturally, our analysis could also be influenced by our specific choices of prior distributions on all model parameters. For instance, does the level of uncertainty about the mutation rate *θ* or the range of growth rates in 𝕂-*exp* change results meaningfully? At least for moderate changes, this seems to be not the case, see A.3.

### Minimal observable clade sizes

We introduced a new statistic, the minimal observable clade size of an individual, defined in Equation (1). It corresponds, for a non-recombining locus, to the size of the smallest genealogical clade this individual is a member of, and that can be distinguished from the (genetic) data. It is not very costly to compute, its computational cost scales well with both population size and mutation rate and, for many scenarios, decreases error rates considerably if (and only if) coupled with other summary statistics. While it needs polarized data it is not taking into account any singleton mutations, which makes it relatively robust towards sequencing errors, as discussed in [Ach08]. Intuitively, the statistic should be suited well for distinguishing between multiple merger genealogies and Kingman’s *n*-coalescent with exponential growth since it is an upper bound of the size of the minimal clade, i.e. the size of the clade spanned by the most recent ancestor node in the genealogy of each individual. For high enough mutation rates, the minimal observable clade size should equal it or be close to it. The minimal clade is known to be typically small for Kingman’s *n*-coalescent, while larger for Beta-*n*-coalescents [BF05], [FSJ14] and [SJY16]. Thus, it would be a well-suited statistic to distinguish the different coalescent models, but it is not directly observable from SNP data (in contrast to the minimal observable clade size). Additionally, the distribution of the minimal clade size is, by definition, invariant under time changes of the coalescent model that do not change its tree topology. Such time-changes of coalescents are the result of non-extreme population size changes in the underlying discrete population models, as mentioned in Section 1.1. This should translate to a weak effect of population size changes on the minimal observable clade sizes. This is true if one compares Kingman’s *n*-coalescent with and without growth, while we see still a positive effect when distinguishing a Dirac *n*-coalescent. However, the latter comparison features lower mutation counts, which makes the minimal and minimal observable clade sizes differ more strongly.

Generally, in all scenarios (when data can be polarized), combining the allele frequency spectrum with statistics from the minimal observable clade sizes produces classification errors for model selection very close to the errors when using all statistics we analyzed. If one further adds five quantiles of the Hamming distances, which may come with a considerable computational burden for larger data sets, one ends up with essentially the same errors as using all statistics. Our recommendation is thus to use this combination for model selection between genealogy models, especially if multiple merger *n*-coalescents are involved.

Interestingly, the harmonic mean of all minimal observable clade sizes often shows one of the highest variable importance among all statistics. A heuristic argument for this is that some of the minimal observable clades may be also minimal clades (i.e. if there is a mutation on the branch adjacent to the ancestor of the minimal clade on its path to the root of the genealogy), thus inheriting the potential of the minimal clade to distinguish between coalescent models. Additionally, not only the information about small clades is included in the harmonic mean. This may prove beneficial, since the sizes of only small minimal observable clades are likely not very informative to distinguish different coalescent models. For instance, the minimal size of all minimal observable clades, at least for large *n* and *θ*, should be 2 for many different coalescent models. For Kingman’s *n*-coalescent, this follows from [DK15, Corollary p.5] and for Beta coalescents from [BBL14, Thm. 8].

### Genealogy model selection confounder population structure

As reported in [KWB19], considering model selection between multiple merger *n*-coalescents and Kingman’s *n*-coalescent with both exponential growth and population structure often causes dramatic misidentification of whether multiple merger coalescents are the correct genealogy models or not. Our results confirm that this is not only a consequence of the use of the singletons and the tail of the site frequency spectrum as the statistics used to perform model selection. Even when adding non-SFS-based statistics, non-accounted population structure makes correct inference of multiple merger genealogies non-feasible.

Can one account for population structure *a priori*? For data sets containing many unlinked loci, assessing population structure may be, at least approximately, possible with standard model-free methods like DAPC [JDB10] or principal component analysis. What about other approaches? Both our results and the results from [KWB19] show that the genetic diversity between model classes with and without population structure (and multiple mergers) is different enough so that additional population structure should be detectable, the open question is how to do it. One potential avenue for (approximate) Bayesian approaches would be to not estimate population structure at all, but instead to distinguish model classes without population structure from a class that combines a genealogy model class with a wide range of population structuring. In other words, one would use a much larger class consisting of a range of prior models, which feature different population structures. We tested this in very limited fashion for populations with two subpopulations of equal size, variable migration rates and subsample sizes with a Kingman’s *n*-coalescent type genealogy with exponential growth, see Section A.4. At least in this scenario, Table A1 shows that Kingman’s *n*-coalescent with exponential growth, Beta-*n*-coalescents and specific models of population structure can be allocated to the true model class with much lower error rates than when just allocating to the models without accounting for population structure. This works both for adding population structure with parameters already featuring in the model class used for training the random forest or not, and does not need a precise *a priori* knowledge or independent (precise) estimation of the population structure.

## 4. Code availability

Please see the public R code repository https://github.com/fabianHOH/mmc_R_gendiv for the code used for coalescent simulation, computation of the diversity statistics and performing the ABC analysis. The repository also contains the ABC results for each model comparison (as `R` objects).

## Acknowledgements

We thank two anonymous referees for suggestions that improved the quality of the paper. FF received funding through DFG grant FR 3633/2-1 through Priority Program 1590: Probabilistic Structures in Evolution. ASJ received funding through CONACyT Grant CB-2014/243068. The authors acknowledge support by the state of Baden-Württemberg through bwHPC.

## Appendix A. Supplementary information

### A.1. Model selection for observed/pseudo-observed data

Given a random forest trained to distinguish between model classes, the approach from [PME^+^15] performs model selection for a set of observed summary statistics by a majority vote. The model that the data is allocated to by the decision trees in the (simple) majority of cases is the selected one. A second approach from [PME^+^15] also allows to give an estimate of the posterior probability for this choice of model *M* : First, for each simulation, the (out-of-bag) misclassifier 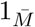 from the majority vote all trees built without them, i.e. 0 if the majority vote is model *M* and 1 otherwise, is recorded. Then, a second random forest from the same simulations is trained to perform a regression of these values 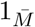 on the summary statistics, i.e. to predict 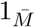 using a random forest of decision trees on the summary statistics (the trees are built slightly different as for model selection, see [PME^+^15] for details), the estimate is then the average over the forest. Then, for the actual observed summary statistic, the posterior probability of classifying as *M* is reported as the average estimate of 1 − 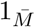 over the second random forest.

We use this approach to assess the effects of model misspecification, i.e. we perform model selection between two or more model classes for simulations from a further, different class. The simulations from the further model class are treated as (pseudo-)observed data: For each simulation, the best fitting model class from the other model classes is recorded (so the winner of the majority vote), alongside its posterior probability.

For this approach, we are not interested in which statistic has the largest potential to distinguish model classes, so we use the non-corrected Gini index to measure how well a statistic at a node of a decision tree sorts the bootstrap samples into model classes.

### A.2. Simulations of robust statistics

For the simulation scheme for Scenario 2 for model classes 𝕂-*exp* and 𝔹, we performed 350,000 simulations of the sets **nSFS, nHam** and **nO** of statistics. Figures A1 to A6 show kernel density estimates of their distributions.

### A.3. Effect on priors on mutation and growth rates

To assess whether changes in mutation prior and range or parameter range affect which statistics distinguish well, we also consider two models who are slightly modificated versions of Scenario 2, named Scenarios 2a and 2b. Sample size is fixed to *n* = 100 and the model selection is performed between 𝕂-*exp* and 𝔹. In Scenario 2a, we fix the number of mutations to *s* = 60 and we consider *ρ ∈* [0.5, 100] plus a prior atom *a* = 0.05 at *ρ* = 0. In Scenario 2b, we state 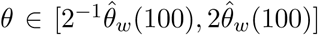 and *ρ* [0.5, 25] with a prior atom *a* = 0.1 at *ρ* = 0. Results are reported in Figures A7 and A8.

### A.4. Using *a priori* arbitrary structured populations to assess multiple mergers vs. exponential growth in structured populations

Assume Scenario 4 with *n* = 100, but replace the fixed migration rate as well as the fixed subsample sizes with scaled migration rates *m* picked uniformly from {0.1, 0.6, … ., 2.1} and subsample sizes picked uniformly from {(10, 90), (20, 80), (30, 70), (60, 40), (50, 50)}. We used the same summary statistics as in the analysis for Scenario 4 in Section 2.4. We generated 175,000 simulations under each scenario twice. Figure A12 shows the result of the three-fold model selection in this scenario, where we denote the model class with population structure and growth by 2*p*-𝕂-*exp*. We then took the random forest trained on the first 175,000 simulations and allocated the set of simulations for model classes 2*p*.5-𝕂-*exp* and 2*p*.9-𝕂-*exp* using ABC based on the constructed random forest and recorded the proportion of misclassification for all simulations and for the ones allocated with high posterior probability as described in Secyion A.1. We then generated a third set of simulations, where we changed migration rates to be uniformly picked from {0.5, 1} and subsample sizes from (5, 95), (25, 75), (45, 55). For these simulations, we denote the model class with population structure and growth by 2*p*\*-𝕂-*exp*. We used the random forest constructed above, allocated also these simulations and recorded the errors as for the second set of simulations. Results are summarized in Table A1.

**Figure A1.**
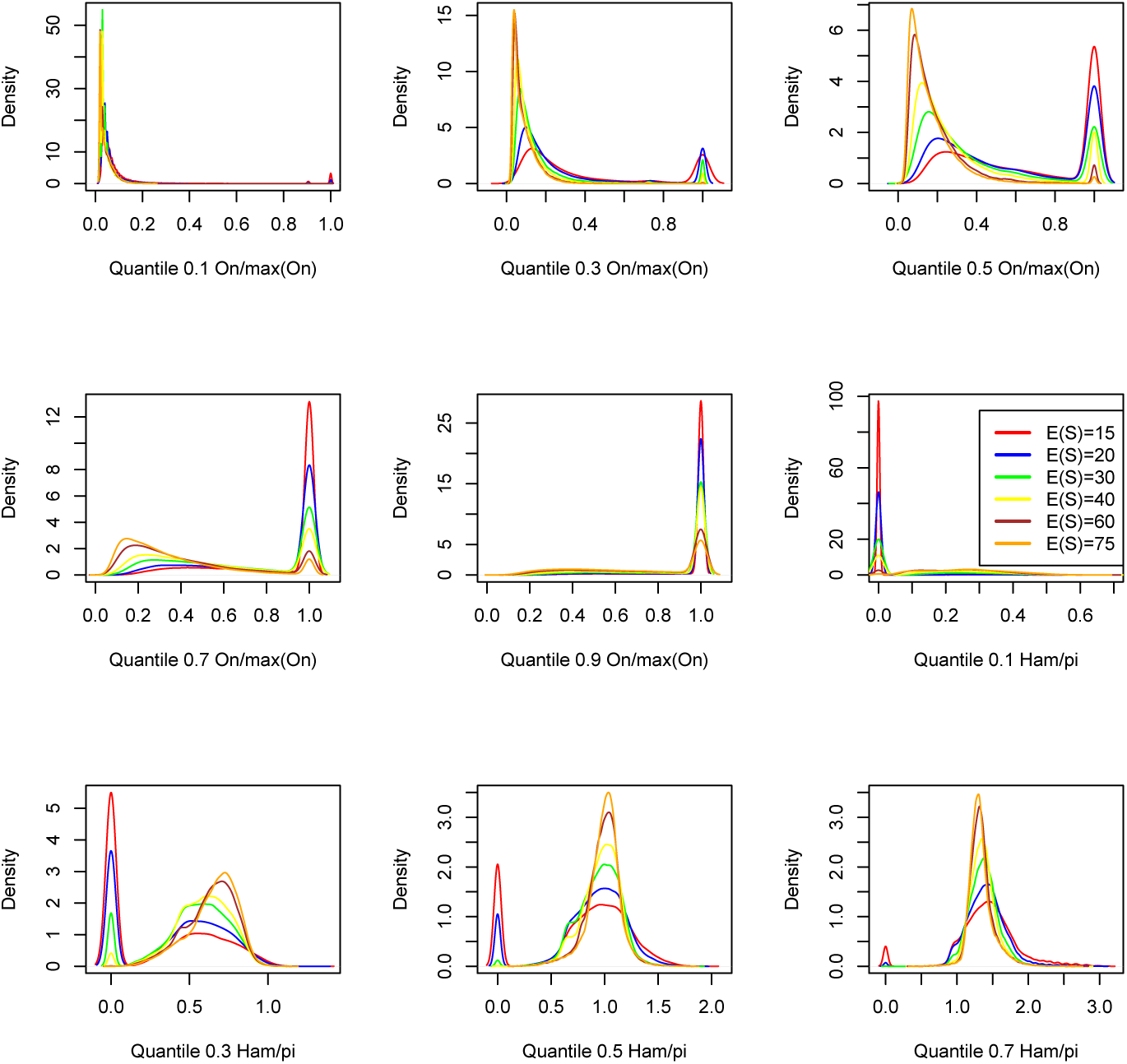
Distributions of statistics **nO** and **nHam** in Scenario 2 for 𝕂-*exp*

### A.5. Comparison with other ABC approaches

For Scenario 2 with *n* = 100 and model classes 𝕂-*exp*, 𝔹 and 𝔻, we performed ABC model selection. We used **O, Ham, Phy, r**^***2***^, **AF**+ as summary statistics with quantiles (.1, .3, …, .9). We compared the ABC with random forest as described in the main manuscript (without bias correction for the variable importances) with three standard ABC approaches as implemented in the R package `abc`: the simple rejection method (REJ), using multinomial logistic regression (MNL) and the neural net approach (NN). For the latter three methods, we used a relatively strict tolerance level of 0.005. All four methods were based on 175,000 simulations per model class as described in the main manuscript. To assess errors, we then performed model selection for two an independent test sets each consisting of 2,500 simulations per model class that then were allocated to the three model classes. For the random forest approach, we proceeded as described in Section A.1, for the other methods we set the model selected to the model with the highest posterior probability. Misclassification is reported in Table A2.

**Figure A2.**
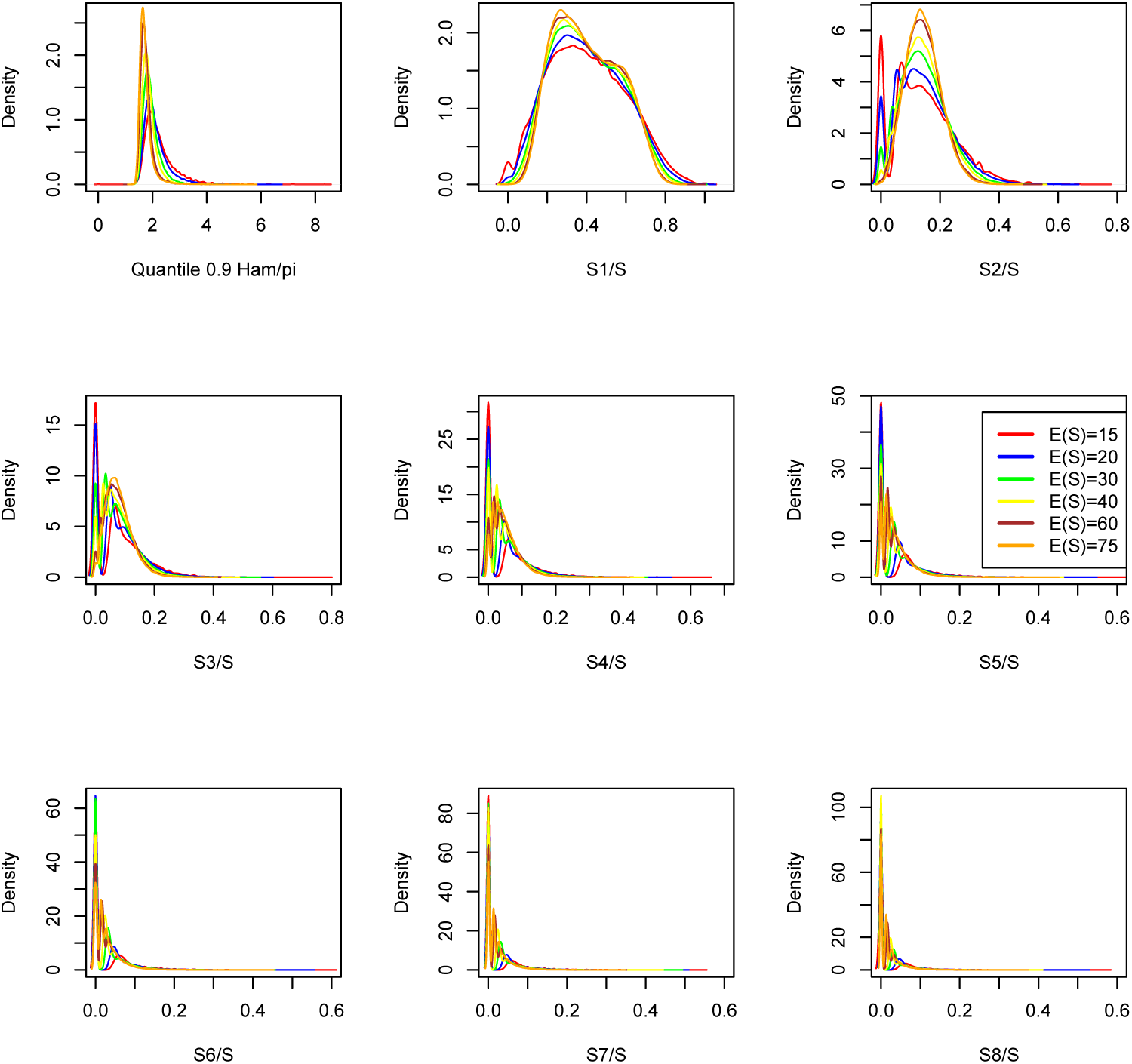
Distributions of statistics **nHam** and **nSFS** in Scenario 2 for 𝕂-*exp*

**Figure A3.**
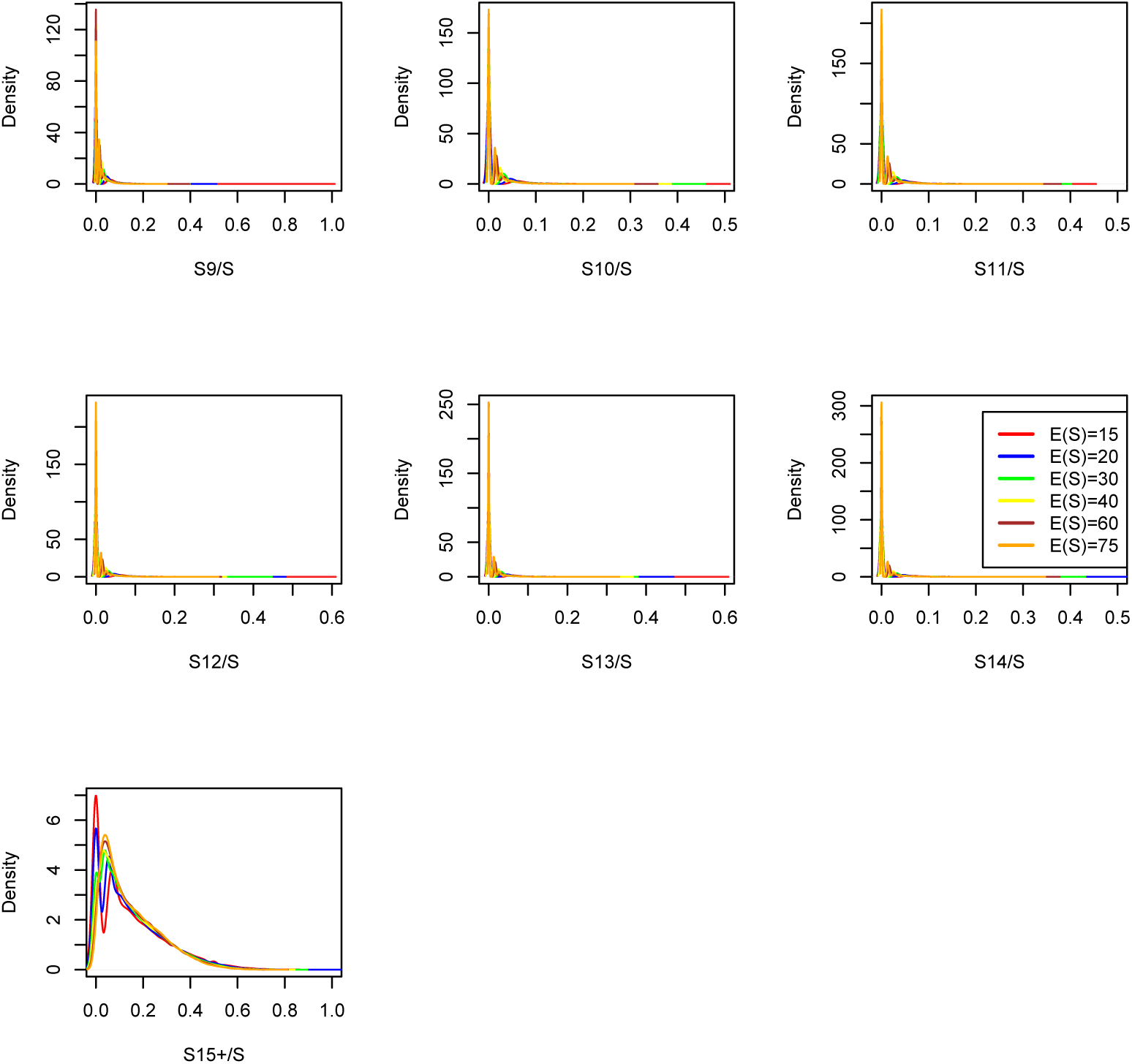
Distributions of statistics **nSFS** in Scenario 2 for 𝕂-*exp*

**Figure A4.**
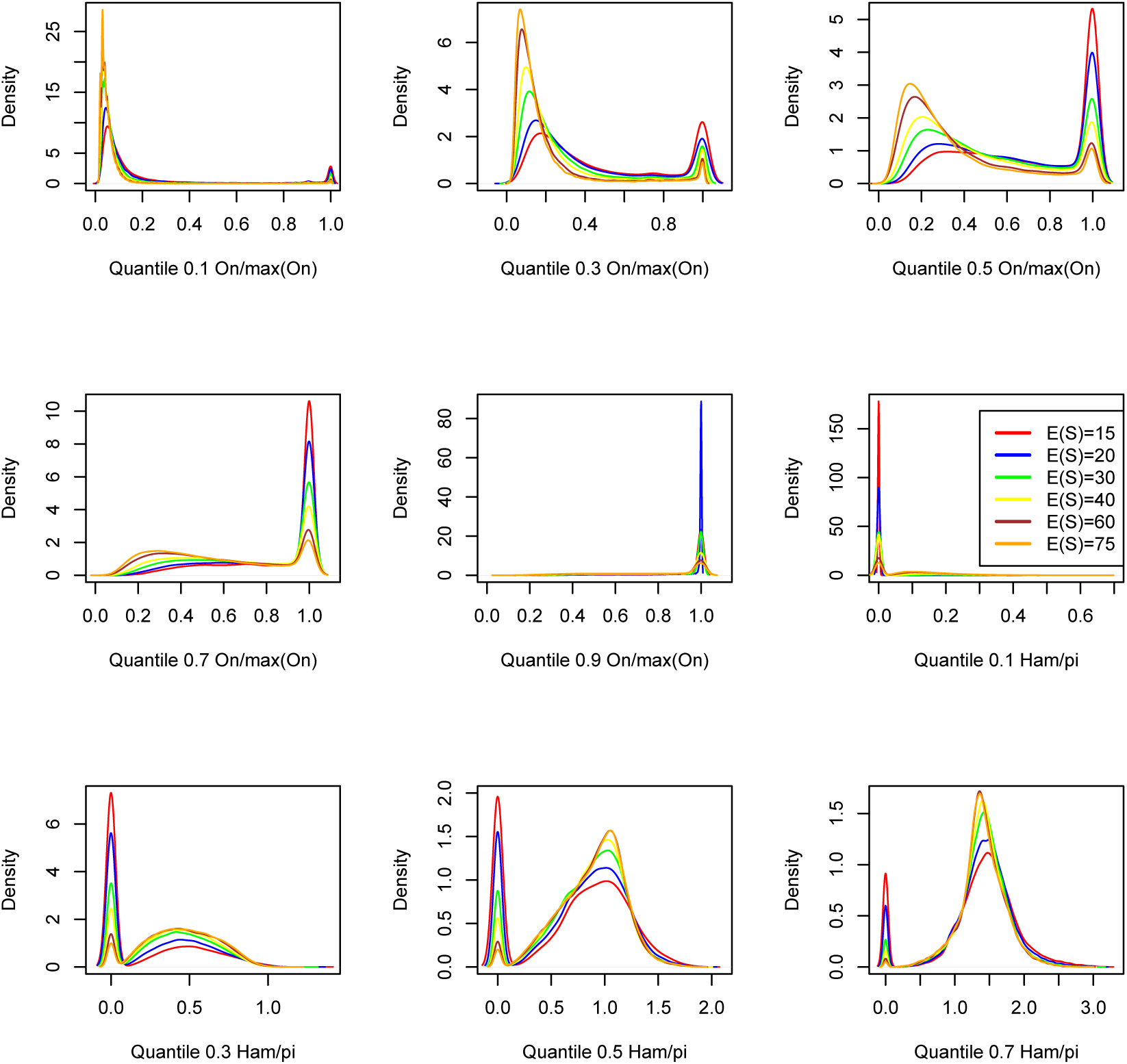
Distributions of statistics **nO** and **nHam** in Scenario 2 for 𝔹

**Figure A5.**
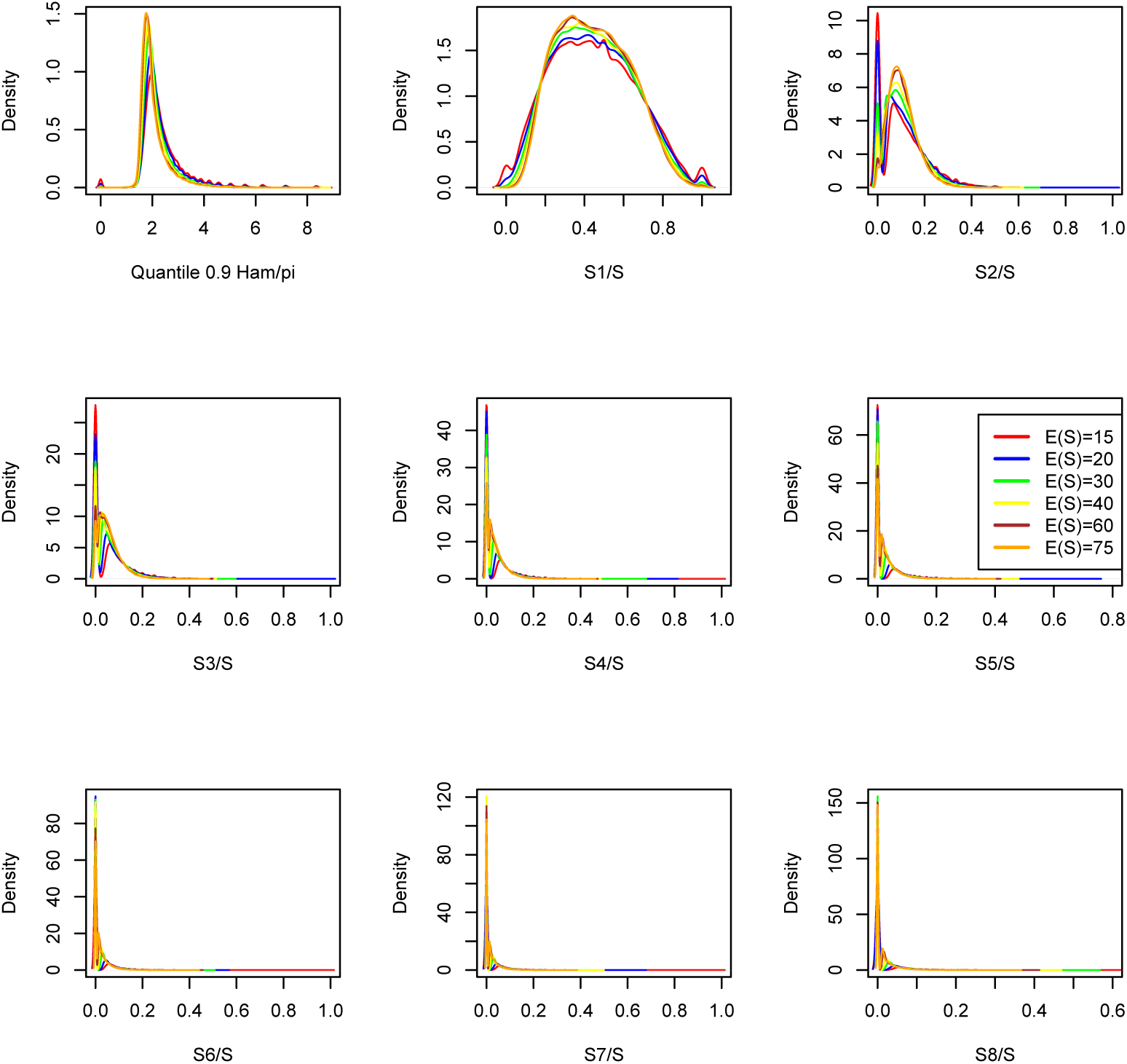
Distributions of statistics **nHam** and **nSFS** in Scenario 2 for 𝔹

**Figure A6.**
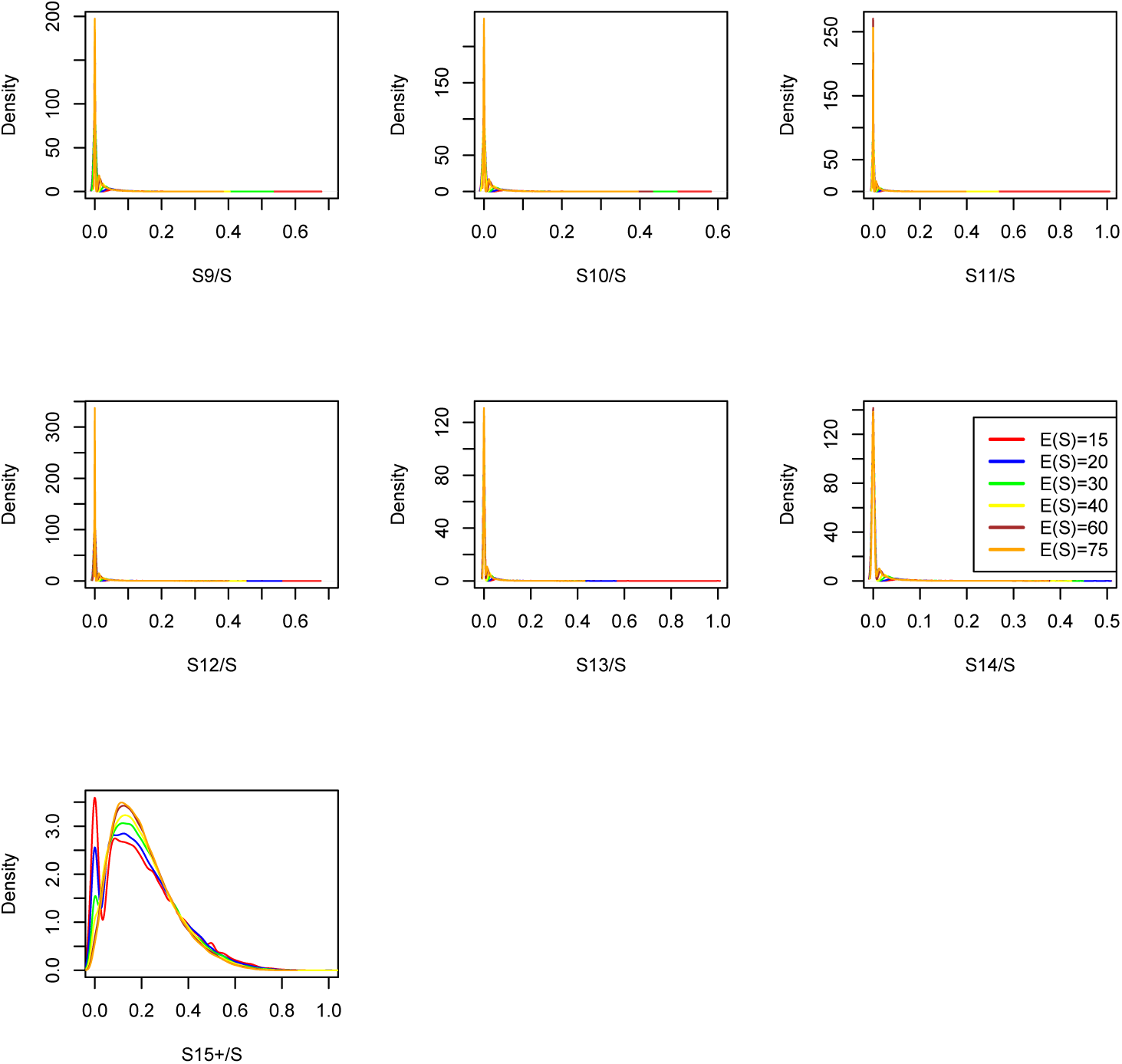
Distributions of statistics **nSFS** in Scenario 2 for 𝔹

**Figure A7.**
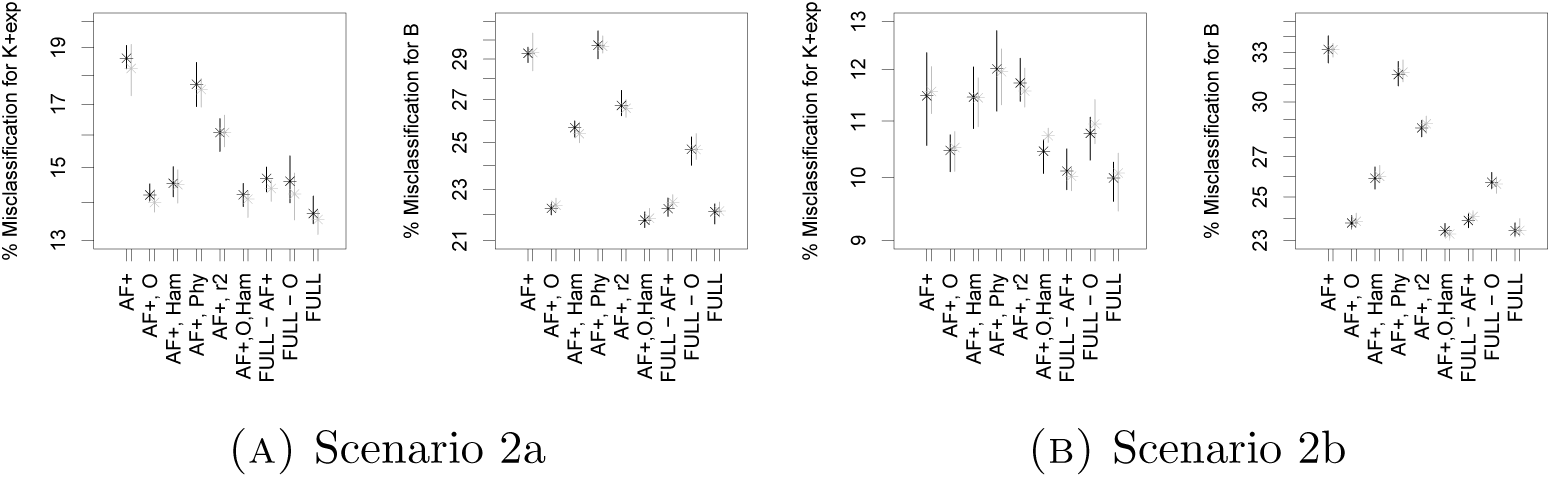
Classification errors (on a log-scale) of model comparison between 𝕂-*exp* and 𝔹 with *n* = 100 for Scenarios 2a and 2b. See Table 7 for the legend. The sets of statistics used are **O, r**^***2***^, **Ham, Phy, AF**+.

**Figure A8.**
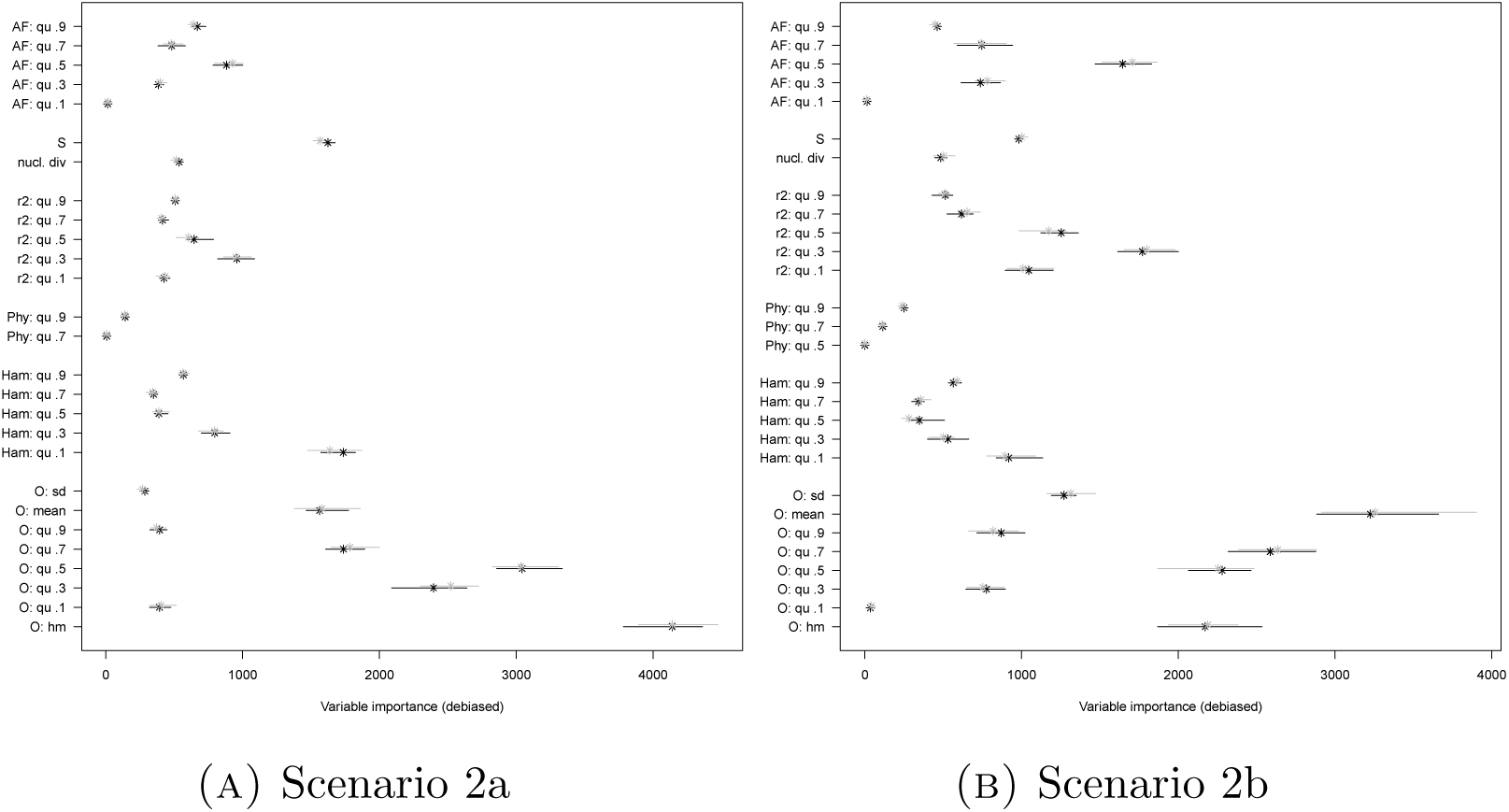
Variable importances for the model comparison between 𝕂-*exp* and 𝔹. See Table 7 for the legend.

**Figure A9.**
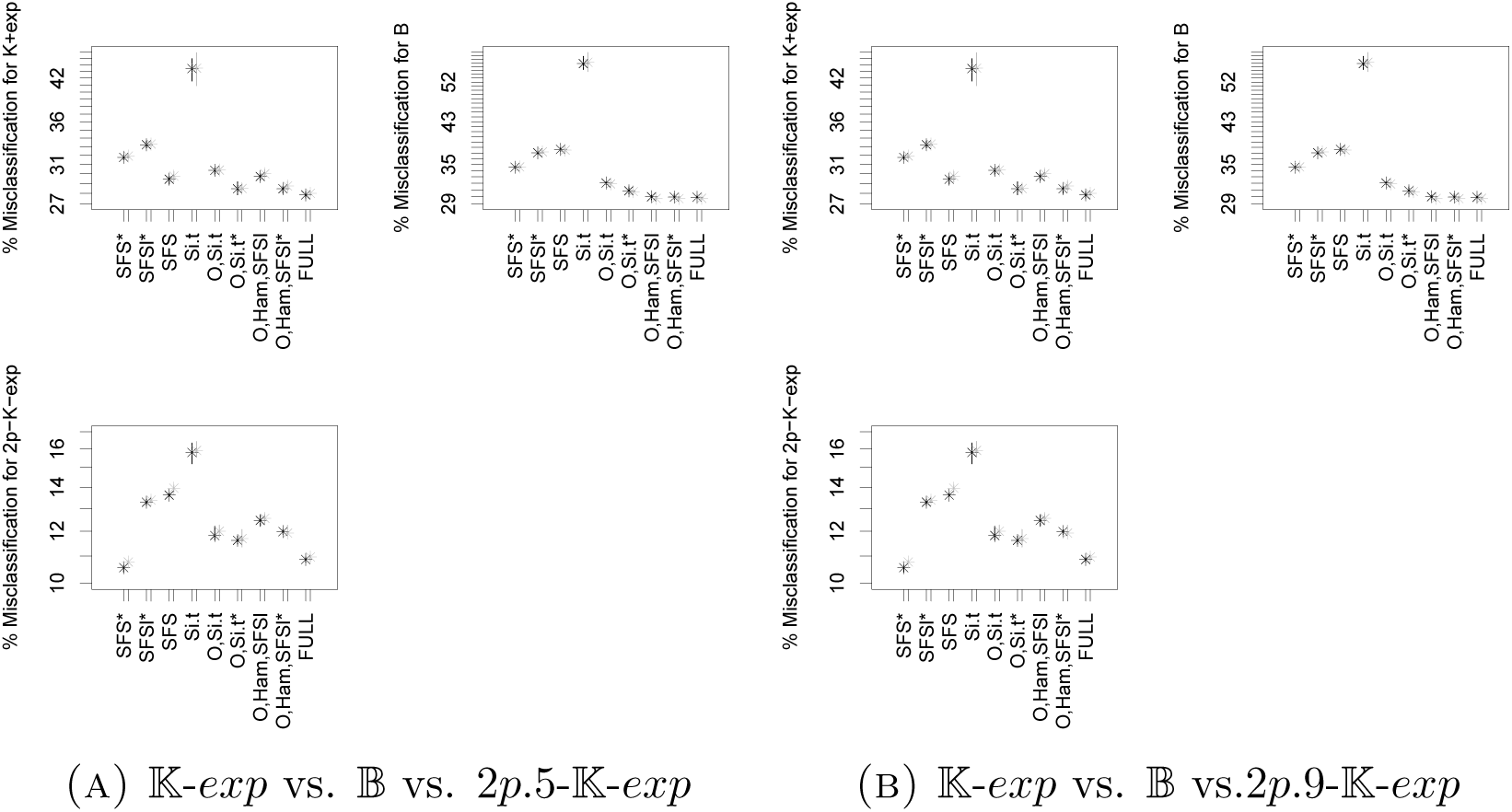
Classification errors (on a log-scale) in Scenario 4, *n* = 100. See Table 7 for the legend. Full set of statistics includes **r**^***2***^, **Phy, O, Ham, SFS**, *π,S*, Tajima’s *D* and Fay and Wu’s *H*.

**Figure A10.**
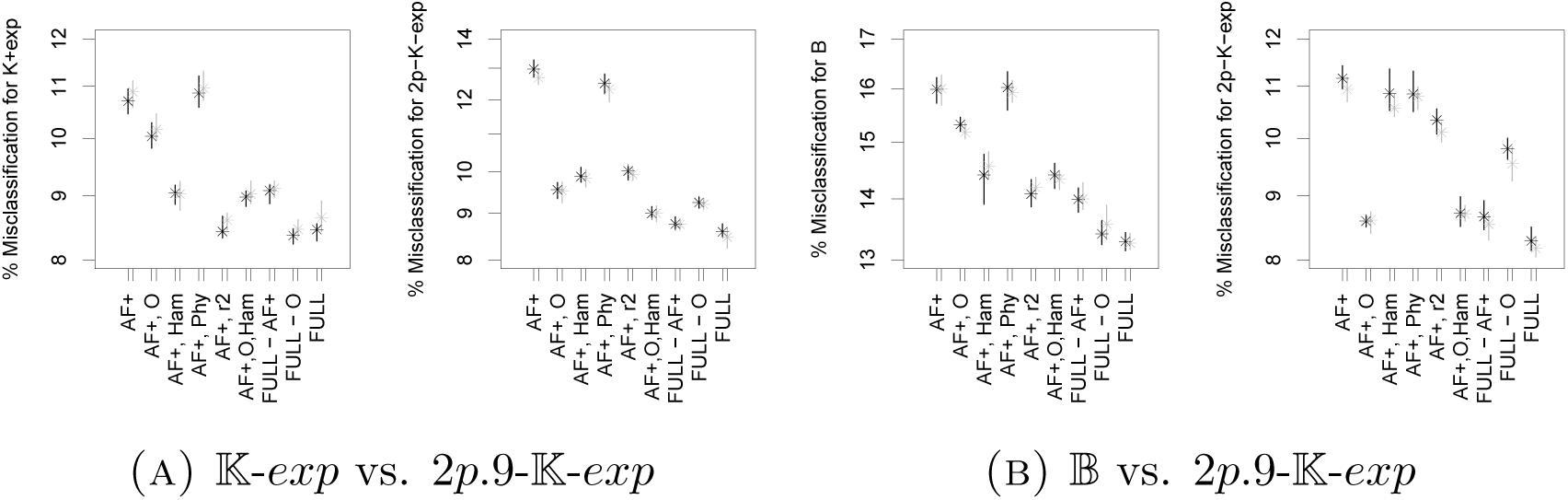
Classification errors (on a log-scale) for model selection in Scenario 4, *n* = 100. See Table 7 for the legend. Full set of statistics includes **r**^***2***^, **Phy, AF**+, **O** and **Ham**.

**Figure A11.**
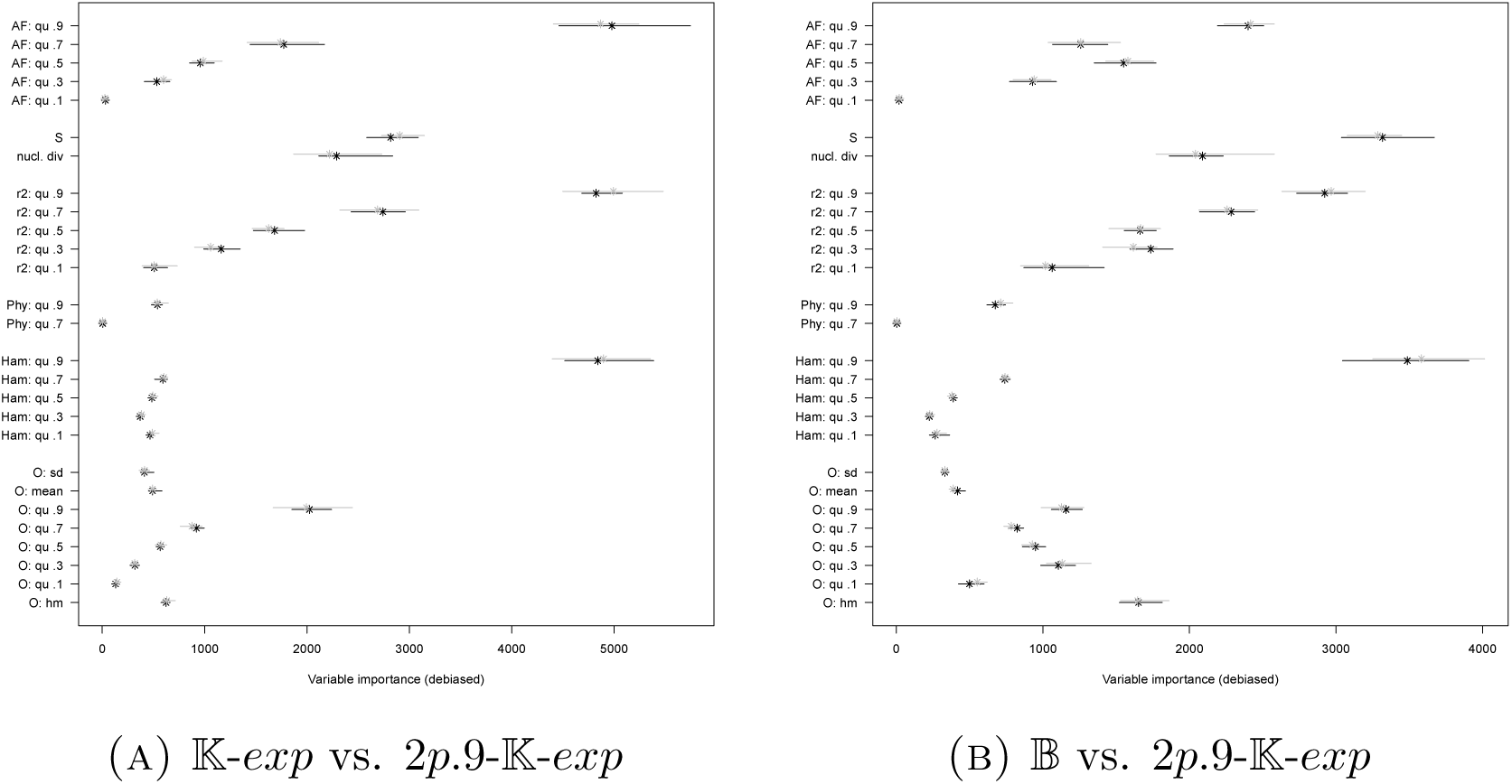
Variable importances in Scenario 4, *n* = 100. See Table 7 for the legend.

**Figure A12.**
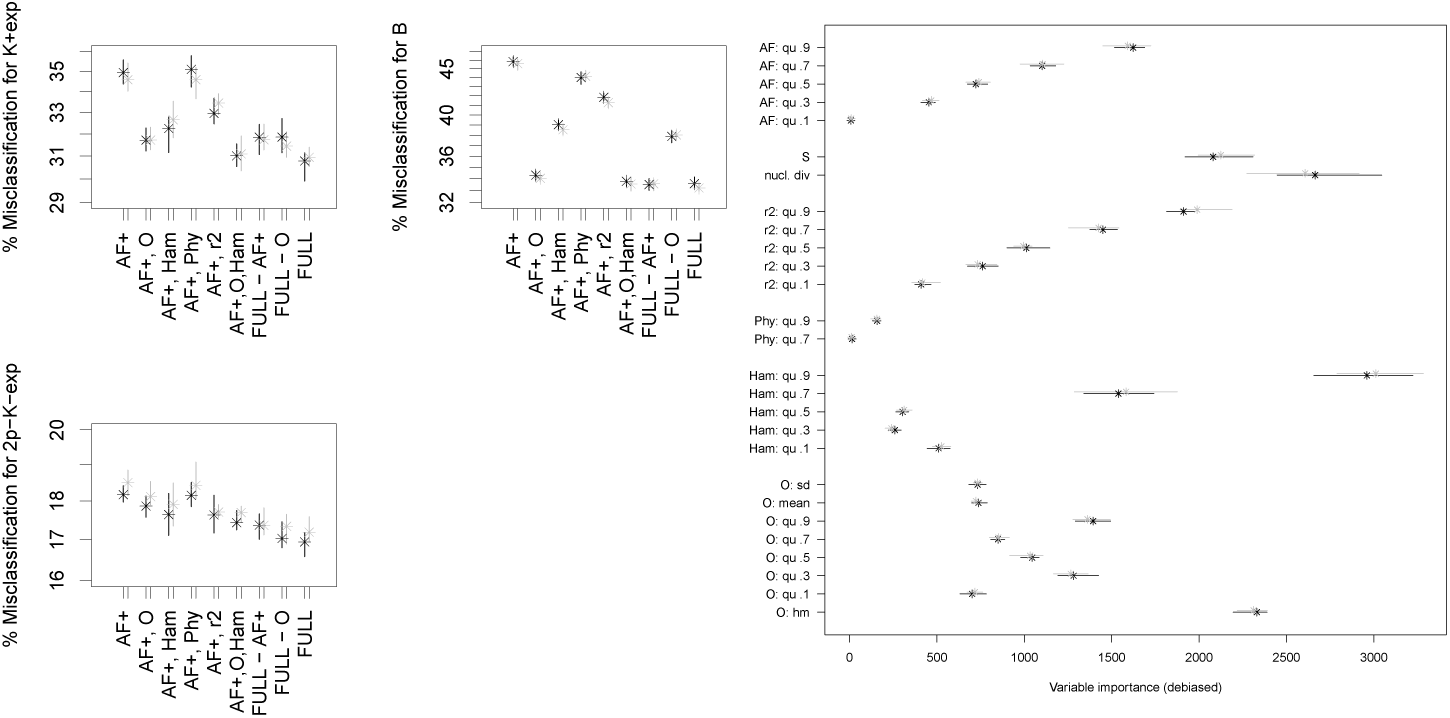
Classification errors (on a log-scale) and variable importances for model selection 𝕂-*exp* vs. 𝔹 vs. 2*p*-𝕂-*exp* (variable migration rates and subsample sizes) in Scenario 4 (*n* = 100). See Table 7 for the legend. Full set of statistics includes **r**^***2***^, **Phy, AF**+, **O** and **Ham**.

**Table A1.** *→ B, → C*: Proportion of simulations from model class *A* allocated to two other model classes *B,C* in Scenarios 2 and 4. In parentheses: same proportion only among high posterior allocations. HP: High posterior allocations are those with posterior probability > 0.9, the column ‘high posterior’ shows their proportion among all simulations from *A*. Random forest based on summary statistics **r**^***2***^, **Phy, SFS***\**, **AF, O** and **Ham** (quantiles used: .1, .2, .3, .4, .9). Results are rounded to two digits, which may cause total allocation probabilities to exceed 1.

| $A$ | $B$ | $C$ | $D$ | $\rightarrow B$ | $\rightarrow C$ | $\rightarrow D$ | HP |
| --- | --- | --- | --- | --- | --- | --- | --- |
| $2p.5\text{-}\mathbb{K}\text{-exp}$ | $\mathbb{K}\text{-exp}$ | $\mathbb{B}$ | $2p\text{-}\mathbb{K}\text{-exp}$ | 0.08 (0) | 0.05 (0) | 0.87 (1) | 0.33 |
| $2p.9\text{-}\mathbb{K}\text{-exp}$ | $\mathbb{K}\text{-exp}$ | $\mathbb{B}$ | $2p\text{-}\mathbb{K}\text{-exp}$ | 0.07 (0.01) | 0.12 (0.01) | 0.81 (0.98) | 0.30 |
| $2p^*\text{-}\mathbb{K}\text{-exp}$ | $\mathbb{K}\text{-exp}$ | $\mathbb{B}$ | $2p\text{-}\mathbb{K}\text{-exp}$ | 0.08 (0.01) | 0.13 (0.04) | 0.80 (0.95) | 0.21 |
| $\mathbb{K}\text{-exp}$ | $\mathbb{K}\text{-exp}$ | $\mathbb{B}$ | $2p\text{-}\mathbb{K}\text{-exp}$ | 0.7 (0.98) | 0.16 (0.02) | 0.14 (0) | 0.09 |
| $\mathbb{B}$ | $\mathbb{K}\text{-exp}$ | $\mathbb{B}$ | $2p\text{-}\mathbb{K}\text{-exp}$ | 0.20 (0.02) | 0.67 (0.97) | 0.13 (0.02) | 0.16 |

**Table A2.**
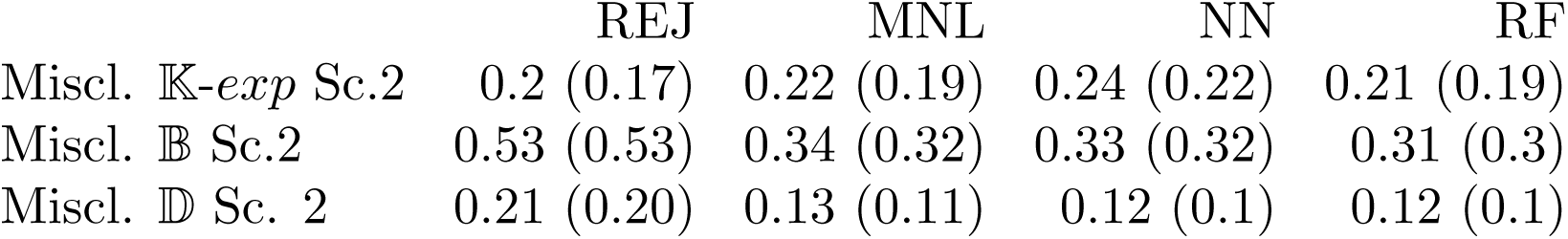
Misclassification frequencies of 2,500 simulations/model class (once replicated, replication result in brackets). ABCs based on MNL=multinomial logistic regression, NN=Neural net, REJ = rejection, RF = random forests.

**Table A3.** Complete mean misidentification matrix (classification errors of model class *A* as model class *B*) of model comparison 𝕂-*exp* vs. 𝔹 vs. 𝔻 vs. 𝔻-*exp* in Scenario 2 with *n* = 100. Result shown for the full set of statistics **r**^***2***^, **Phy, Ham, O** and **AF**+. Results averaged over 2× 10 ABC runs from each of two simulation sets and rounded to three digits.

| | as $\mathbb{K}\text{-exp}$ | as $\mathbb{B}$ | as $\mathbb{D}$ | as $\mathbb{D}\text{-exp}$ |
| --- | --- | --- | --- | --- |
| from $\mathbb{K}\text{-exp}$ | 0.802 | 0.191 | 0.001 | 0.006 |
| from $\mathbb{B}$ | 0.246 | 0.680 | 0.047 | 0.027 |
| from $\mathbb{D}$ | 0.014 | 0.098 | 0.534 | 0.354 |
| from $\mathbb{D}\text{-exp}$ | 0.012 | 0.049 | 0.192 | 0.747 |

